# Compressed Perturb-seq: highly efficient screens for regulatory circuits using random composite perturbations

**DOI:** 10.1101/2023.01.23.525200

**Authors:** Douglas Yao, Loic Binan, Jon Bezney, Brooke Simonton, Jahanara Freedman, Chris J. Frangieh, Kushal Dey, Kathryn Geiger-Schuller, Basak Eraslan, Alexander Gusev, Aviv Regev, Brian Cleary

**Affiliations:** Program in Systems, Synthetic, and Quantitative Biology, Harvard University, Cambridge, MA; Klarman Cell Observatory, Broad Institute of Harvard and MIT, Cambridge, MA; Department of Electrical Engineering and Computer Science, Massachusetts Institute of Technology, Cambridge, MA; Harvard T.H. Chan School of Public Health, Boston, MA; Genentech, South San Francisco, CA; Department of Medical Oncology, Dana-Farber Cancer Institute, Boston, MA; Division of Genetics, Brigham and Women’s Hospital, Boston, MA; Faculty of Computing and Data Sciences, Boston University, Boston, MA; Department of Biology, Boston University, Boston, MA; Department of Biomedical Engineering, Boston University, Boston, MA; Program in Bioinformatics, Boston University, Boston, MA; Biological Design Center, Boston University, Boston, MA; Department of Genetics, Stanford University School of Medicine, Stanford, CA

## Abstract

Pooled CRISPR screens with single-cell RNA-seq readout (Perturb-seq) have emerged as a key technique in functional genomics, but are limited in scale by cost and combinatorial complexity. Here, we reimagine Perturb-seq’s design through the lens of algorithms applied to random, low-dimensional observations. We present compressed Perturb-seq, which measures multiple random perturbations per cell or multiple cells per droplet and computationally decompresses these measurements by leveraging the sparse structure of regulatory circuits. Applied to 598 genes in the immune response to bacterial lipopolysaccharide, compressed Perturb-seq achieves the same accuracy as conventional Perturb-seq at 4 to 20-fold reduced cost, with greater power to learn genetic interactions. We identify known and novel regulators of immune responses and uncover evolutionarily constrained genes with downstream targets enriched for immune disease heritability, including many missed by existing GWAS or trans-eQTL studies. Our framework enables new scales of interrogation for a foundational method in functional genomics.

## INTRODUCTION

Pooled perturbation screens with high-content readouts, ranging from single-cell RNA-seq (Perturb-seq)^1–4^ to imaging-based spatial profiling^5–7^, are now enabling systematic studies of the regulatory circuits that underlie diverse cell phenotypes. Perturb-seq has been applied to various model systems, leading to insights about diverse cellular processes including the innate immune response^2^, *in vivo* effects of autism risk genes in mice^8^ and organoids ^9,10^, and genome-scale effects on aneuploidy, differentiation, and RNA splicing^11^. Integrating data from population-level genetic screens has also elucidated human disease mechanisms^12^.

However, due to the large number of genes in the genome, large-scale Perturb-seq screens are still prohibitively expensive and are often limited by the number of available cells, especially for primary cell systems^13^ and *in vivo* niches^8^. In addition, the exponentially larger number of possible genetic interactions makes it impossible to conduct exhaustive combinatorial screens for genetic interactions using existing approaches, so current Perturb-seq studies of genetic interactions are very modest and focused^14^. Several approaches have been developed to improve the efficiency of scRNA-seq and/or Perturb-seq, including overloading droplets with multiple pre-indexed cells (SciFi-seq^15^) or pooling multiple guides within cells^16^. However, pre-indexing requires an additional laborious and complex experimental step, while guide pooling has only been used to study *cis* and not *trans* effects of perturbations.

We propose an alternative approach to greatly increase the efficiency and power of Perturb-seq for both single and combinatorial perturbation screens, inspired by theoretical results from compressed sensing^17–19^ that apply to the sparse and modular nature of regulatory circuits in cells. To elaborate, perturbation effects tend to be *sparse* in that most perturbations affect only a small number of genes or co-regulated gene programs^2^. In this scenario, rather than assaying each perturbation individually, we can measure a much smaller number of *random combinations* of perturbations (forming what we call “composite samples”) and accurately learn the effects of individual perturbations from the composite samples using sparsity-promoting algorithms. Moreover, with certain types of composite samples, we can efficiently learn both first-order effects (*i*.*e*., from single gene perturbations) and higher-order genetic interaction effects from the same data. We have previously shown that experiments that measure random compositions of the underlying biological dataset can greatly increase the efficiency of measuring expression profiles^20^ and imaging transcriptomics^21^.

Here, we develop two experimental strategies to generate composite samples for Perturb-seq screens, and we introduce an inference method, Factorize-Recover for Perturb-seq analysis (FR-Perturb), to learn individual perturbation effects from composite samples. We apply our approach to 598 genes in a human macrophage cell line treated with lipopolysaccharide (LPS). By comparing compressed Perturb-seq to conventional Perturb-seq conducted in the same system, we demonstrate the enhanced efficiency and power of our approach for learning single perturbation effects and second-order genetic interactions. We derive insights into immune regulatory functions and illustrate their connection to human disease mechanisms by integrating data from genome-wide association studies (GWAS) and expression quantitative trait loci (eQTL) studies.

## RESULTS

### A compressed sensing framework for perturbation screens

In conventional Perturb-seq, each cell in a pool receives one or more genetic perturbations. Each cell is then profiled for the identity of the perturbation(s) and the expression levels of *m* ≈ 20,000 expressed genes. Our goal is to infer the effect sizes of *n* perturbations on the phenotype, which can be the entire gene expression profile (*n* × *m* matrix) or an aggregate multi-gene phenotype^2,3,11^ such as an expression program or cell state score (length-*n* vector). In both cases, we need *O*(*n*) samples to learn the effects of *n* perturbations (**Figure 1A**) (where sample replicates introduce a constant factor that is subsumed under the big O notation), such that the number of samples scales *linearly* with the number of perturbations.

**Figure 1.**
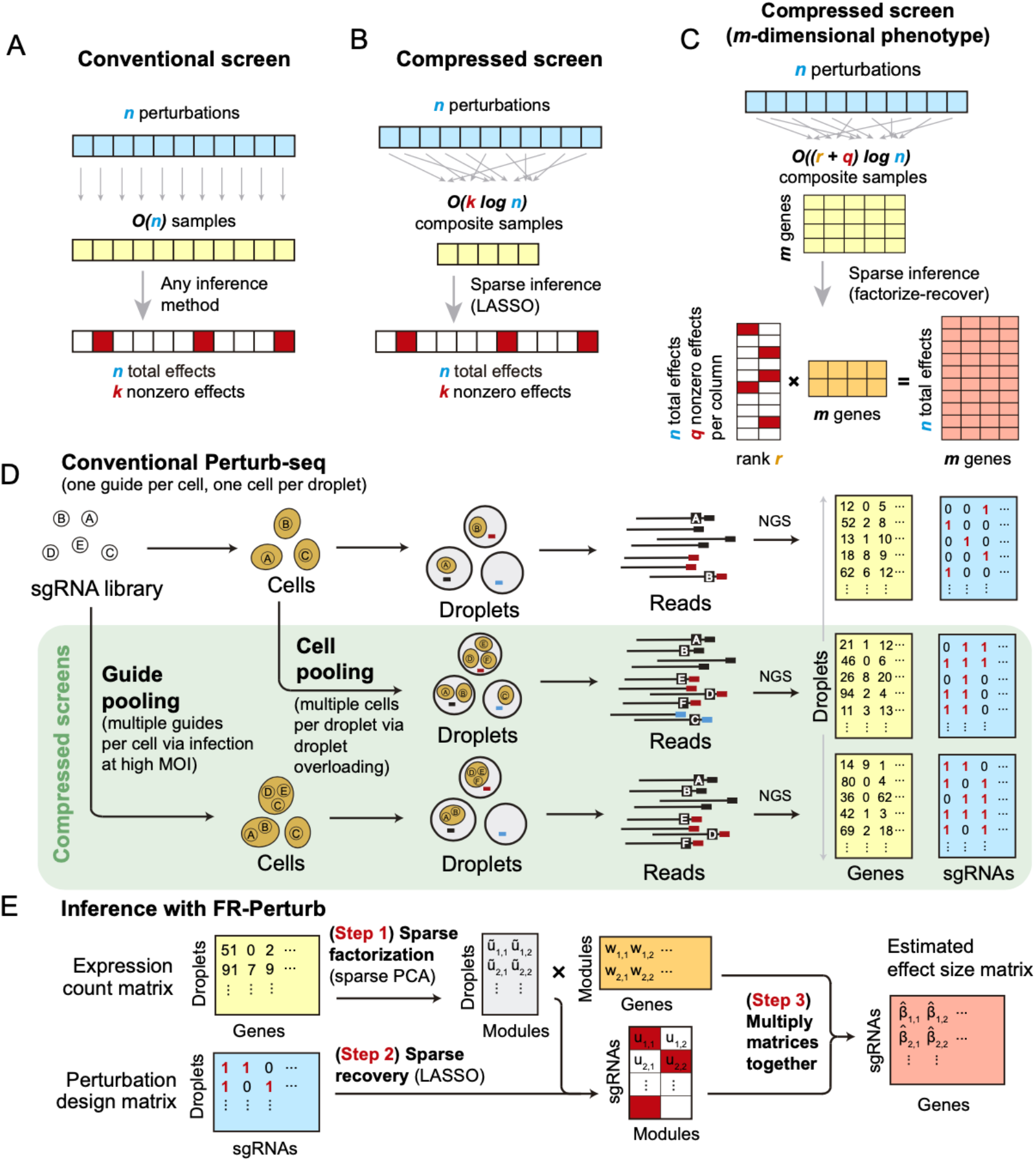
Framework for compressed Perturb-seq. (**A**) Schematic for conventional perturbation screen with single-valued phenotype. Each sample (yellow) receives a single perturbation (blue). The required number of samples scales linearly with the number of perturbations, as captured by the *O*(*n*) term. (**B**) Schematic for compressed perturbation screen with single-valued phenotype. Each “composite” sample (yellow) represents a random combination of perturbations (blue). The required number of samples scales sub-linearly with the number of perturbations given the following: (1) the effects of the perturbations are sparse (*i*.*e*., *k* increases more slowly than *n*), and (2) sparse inference (typically LASSO) is used to infer the effects from the composite sample phenotypes. (**C**) Schematic for compressed perturbation screen with high-dimensional phenotype, which is the main use case for Perturb-seq. The required number of samples scales sub-linearly with the number of perturbations given the following: (1) the effects of the perturbations are sparse and act on a relatively small number of groups of correlated genes (*i*.*e*., *q* and *r* increase more slowly than *n*), and (2) sparse inference (namely the “factorize-recover” algorithm^22^) is used to infer the effects from the composite sample phenotypes. (**D**) Two experimental strategies for generating composite samples for Perturb-seq. Both “cell pooling” and “guide pooling” change one step of the conventional Perturb-seq protocol. The result is a sample whose phenotype corresponds to a random linear combination of the phenotypes of samples from the conventional Perturb-seq screen. (**E**) Schematic of computational method used to infer perturbation effects from composite sample phenotypes, based on the “factorize-recover” algorithm^22^.

Based on the theory of compressed sensing^17^, there exist conditions under which far fewer than *O*(*n*) samples are sufficient to learn the effects of *n* perturbations. In general, if the perturbation effects are sparse (*i*.*e*., relatively few perturbations affect the phenotype), or are sparse in a latent representation (*i*.*e*., perturbations tend to affect relatively few latent factors that can be combined to “explain” the phenotype), then we can measure a small number of random composite samples (comprising *linear combinations* of individual sample phenotypes) and decompress those measurements to infer the effects of individual perturbations. Composite samples can be generated either by randomly pooling perturbations in individual cells, or by randomly pooling cells containing one perturbation each (see below).

The number of required composite samples depends on whether the phenotype is single-valued or high-dimensional. When the phenotype is single-valued (*e*.*g*., fitness), *O*(*k* log *n*) composite samples suffice to accurately recover the effects of *n* perturbations^18,19^, where *k* is the number of nonzero elements among the *n* perturbation effects (**Figure 1B**). When most genes do not affect the phenotype, *k* grows more slowly than *n*, and the number of required composite samples scales logarithmically or at worst sub-linearly with the number of perturbations. Meanwhile, when the phenotype is an *m*-dimensional gene expression profile, an efficient approach involves inferring effects on latent expression factors, then reconstructing the effects on individual genes from these factors using the “factorize-recover” algorithm^22^. This approach requires *O*((*q* + *r*) log*n*) composite samples, where *r* is the *rank* of the *n* × *m* perturbation effect size matrix (*i*.*e*., the maximum number of its linearly independent column vectors), and *q* is the maximum number of nonzero elements in any column of the left matrix of the factorized effect size matrix (**Figure 1C**). In our case, *r* is the number of distinct groups of “co-regulated” genes whose expression changes concordantly in response to any perturbation, while *q* is the maximum number of “co-functional” perturbations with nonzero effects on any individual module. Due to the modular nature of gene regulation^23,24^, *r* and *q* are expected to remain small when *n* increases. Indeed, we observed a relatively small number of co-functional and co-regulated gene groups (small *q* and *r*, respectively, relative to *n*) in previous Perturb-seq screens in various systems^2,13^. Thus, the number of required composite samples will scale logarithmically or at worst sub-linearly with *n*, leading to much fewer required samples than the conventional approach with large *n*.

### Experimentally generating composite samples for compressed Perturb-seq

We generated composite samples for compressed Perturb-seq by either randomly pooling cells containing one perturbation each in overloaded scRNA-seq droplets^15^, or by randomly pooling guides in individual cells via infection with a high multiplicity of infection (MOI)^2,16^ (**Figure 1D**). Either method can be used to learn the same underlying perturbation effects, but each has different strengths and limitations (**Discussion**).

For pooling cells with droplet overloading, we load cells into the microfluidics chip at much higher concentration than usual so that each droplet contains multiple cells. The resulting expression counts from a given droplet are proportional to the average expression counts of the cells in the droplet, while the set of detected guides is the union of those present in any of the constituent cells. We do not intend to match reads to specific cells within each droplet and in fact cannot do so without additional indexing^15^. Rather, we model the expression counts of each gene as the expected counts from an unperturbed (control) cell multiplied by the average of the fold-change effects from each guide in the droplet (**Methods**).

For pooling guides, we transduce cells at high MOI so that multiple random guides enter each cell. We then measure the expression counts from individual cells and identify their constituent guides. We model the log-expression counts of each gene as the log of the expected counts from a control cell plus the sum of log fold-change effects from the guides in the cell (**Methods**). This model, where observations are a random linear combination of log fold-change effect sizes from different guides, makes the non-trivial assumption that the effect sizes of guides tend to combine additively in log expression space when present in the same cell. We can test the validity of this assumption by looking at the concordance between effect size estimates from cells with only one guide versus cells with multiple guides (see below). Pooling guides has a key benefit over pooling cells, in that the generated data can be used to estimate both first-order effects and higher-order genetic interactions (with appropriate sample sizes and explicit interaction terms in the model) (**Methods**). We illustrate the feasibility of estimating second-order effects from our guide-pooled data below.

### FR-Perturb infers effects from compressed Perturb-seq

To infer perturbation effects from the composite samples, we devised a method called FR-Perturb based on the “factorize-recover” algorithm^22^ (**Methods**). FR-Perturb first factorizes the expression count matrix with sparse factorization (*i*.*e*., sparse PCA), followed by sparse recovery (*i*.*e*., LASSO) on the resulting left factor matrix comprising perturbation effects on the latent factors. Finally, it computes perturbation effects on individual genes as the product of the left factor matrix from the recovery step with the right factor matrix (comprising gene weights in each latent factor) from the first factorization step (**Figure 1E**; **Methods**). We obtained p-values and false discovery rates (FDR) for all effects by permutation testing (**Methods**). We evaluated FR-Perturb by comparing it to existing inference methods for Perturb-seq, namely elastic net regression^2^ and negative binomial regression^16^ (see below).

### Compressed Perturb-seq screens of the LPS response

We implemented and evaluated compressed Perturb-seq in the response of THP1 cells (a human monocytic leukemia cell line) to stimulation with LPS when either pooling cells or pooling guides (**Figure 2A-B**). In each case, we also performed conventional Perturb-seq, targeting the same genes in the same system for comparison. We selected 598 genes to be perturbed from seven mostly non-overlapping immune response studies (**Table S1**), including genes with roles in the canonical LPS response pathway (34 genes), GWAS for inflammatory bowel disease (79 genes) and infection (106 genes), Mendelian immune diseases from OMIM with keywords for “bacterial infection” (85 genes) and “NF-kappa-B” (102 genes), a previous genome-wide screen for effects on TNF expression in mouse BMDCs^25^ (93 genes), and genes with large genetic effects in *trans* on gene expression from an eQTL study in patient-derived macrophages stimulated with LPS^26^ (79 genes) (**Methods, Figure S1A**). We designed 4 sgRNAs for each gene and 500 each of non-targeting or safe-targeting control sgRNAs, resulting in a total pool of 3,392 sgRNAs (**Methods**). We introduced the sgRNAs into THP1 cells via a modified CROP-seq vector^4^ (**Methods**). After transduction and selection, we treated cells with PMA for 24 hours and grew them for another 48 hours as they differentiated into a macrophage-like state^27^, then treated them with LPS for three hours before harvesting for scRNA-Seq (**Methods**). For our cell-pooled screen, we used CRISPR-Cas9 to knock out genes^2^, whereas for our guide-pooled screen we used CRISPRi with dCas9-KRAB to knock down gene expression^1^ (**Figure 2A**) to avoid cellular toxicity due to multiple double-stranded breaks in individual cells^28^.

**Figure 2.**
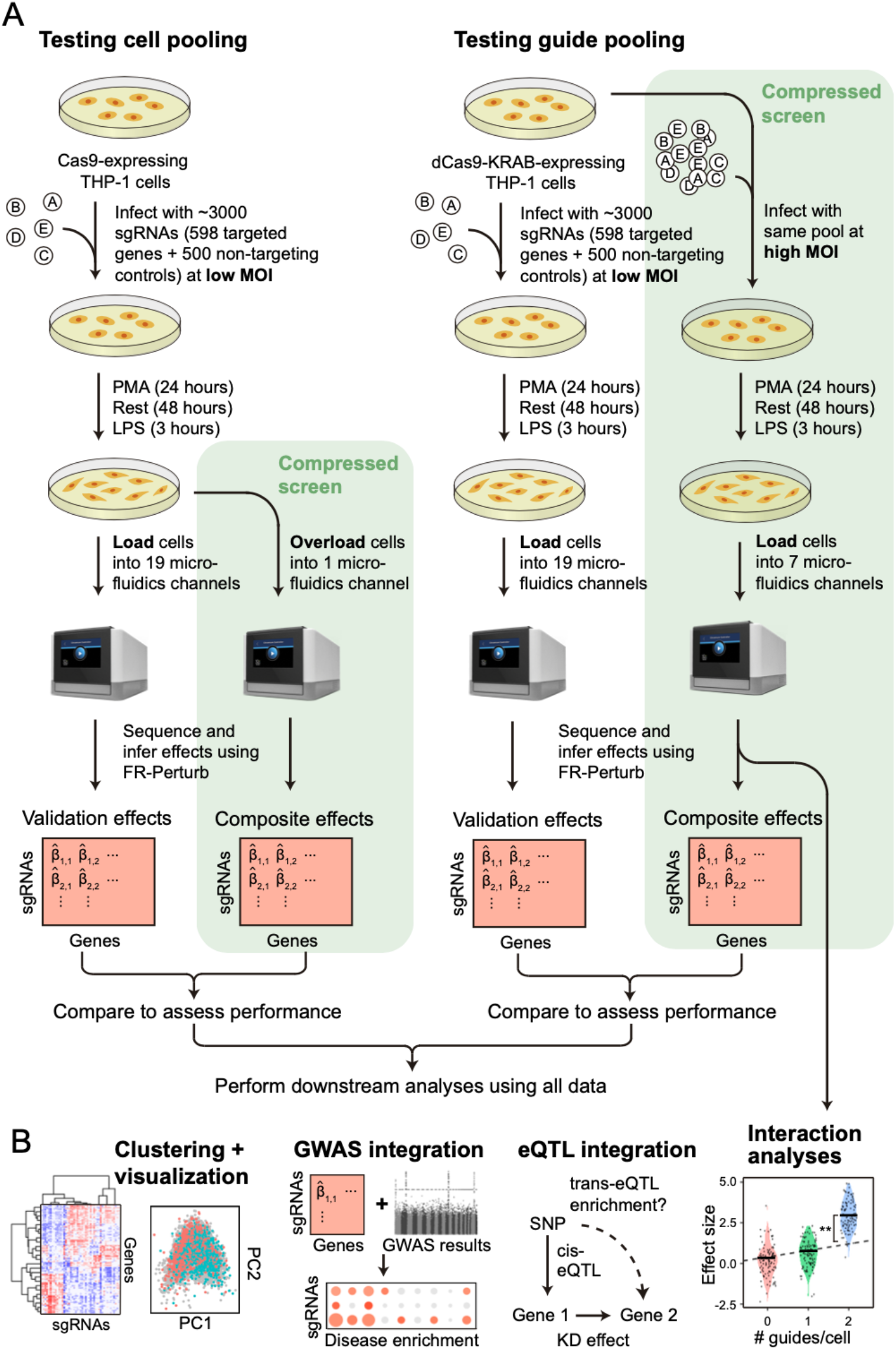
Experimental overview. (**A**) Outline of experiments used to test and validate cell pooling (left) and guide pooling (right). (**B**) Downstream analyses performed using perturbation effects from all experiments.

By design, the two compressed screens were substantially smaller than their corresponding conventional screens. In the cell-pooled screen, we analyzed a single channel of droplets (10x Genomics, **Methods**) overloaded with 250,000 cells, while for the corresponding conventional Perturb-seq screen we analyzed 19 channels at normal loading. We sequenced the library from the overloaded channel to a depth of 4-fold more reads than a conventional channel to account for the larger number of non-empty droplets and greater expected RNA content per droplet. After quality control, there were 32,700 droplets containing at least one sgRNA from the overloaded channel (*vs*. 4,576 droplets/channel for a total of 86,954 droplets from the conventional screen) (**Figure 3A**), with a mean of 1.86 sgRNAs per non-empty droplet (conventional: 1.11) (**Figure 3B**) and a mean of 90 droplets containing a guide for each perturbed gene (conventional: 144) (**Figure 3C**). We observed 14,987 total genes with measured expression (conventional: 17,552). Thus, the cell-pooled screen had >7 times the number of non-empty droplets per channel compared to the conventional screen; considering library preparation and sequencing costs, it was approximately 8 times cheaper.

**Figure 3.**
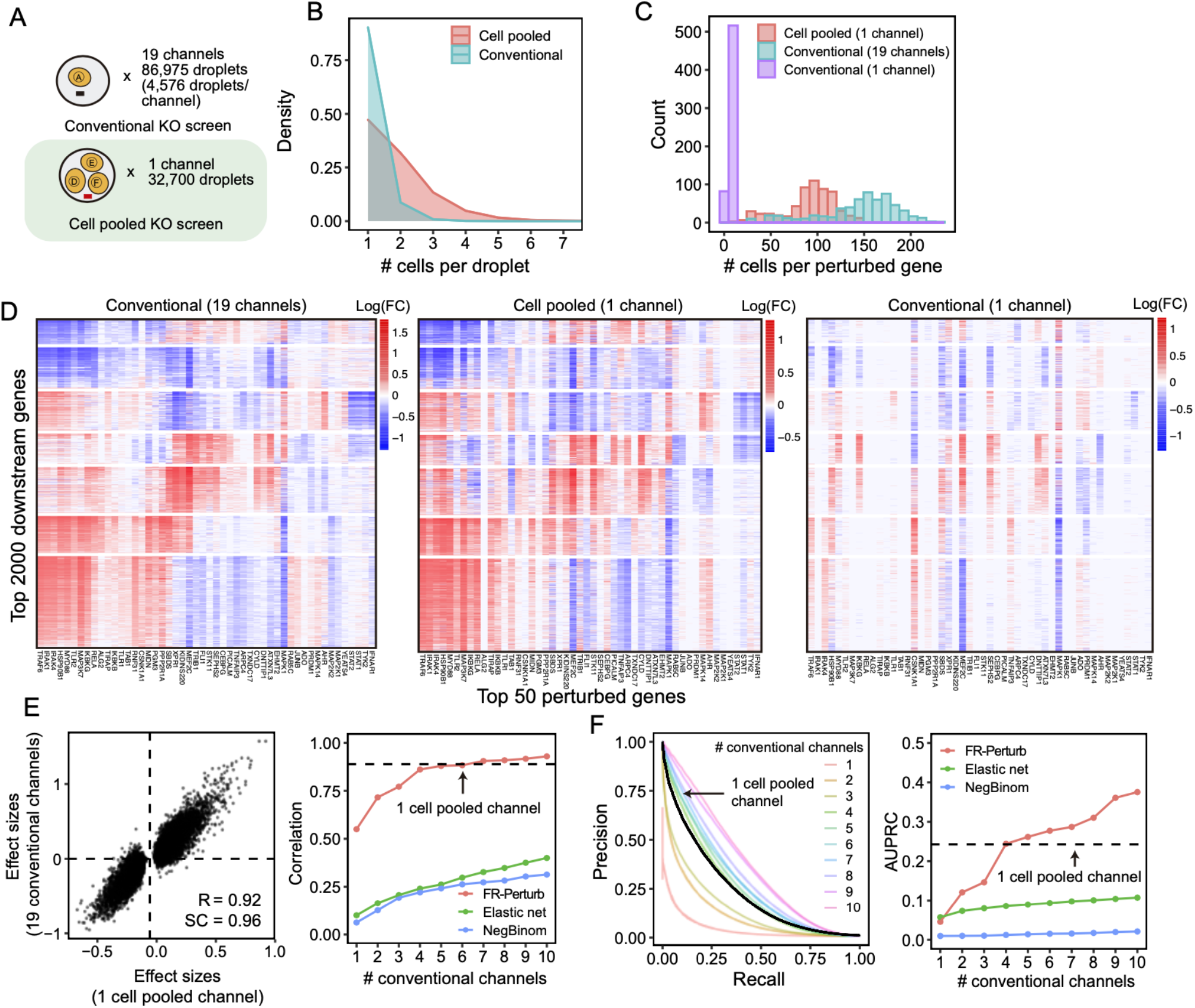
Evaluating cell-pooled Perturb-seq versus conventional Perturb-seq. (**A**) Number of channels and droplets from the conventional validation screen (top) and cell-pooled screen (bottom). (**B**) Distribution of droplets based on number of cells they contain for the cell-pooled and conventional screens. In practice, we only directly measure the number of guides/droplet rather than cells/droplets, but these quantities are equivalent given 1 guide/cell. (**C**) Distribution of the number of cells containing a guide targeting each perturbed gene in the cell-pooled screen and conventional screen. For the conventional screen, we show the distributions for both the full screen (19 channels) and an equivalent number of channels as the cell-pooled screen (1 channel). (**D**) Heatmaps of the top effect sizes (inferred with FR-Perturb) from the conventional screen (left), with the same effect sizes shown for the cell pooled screen (middle) and one equivalent channel of the conventional screen (right). X-axis: top 50 perturbed genes, based on their average magnitude of effect on all 17,552 downstream genes. Y-axis: top 2,000 downstream genes, based on the average magnitude of effects of all 598 perturbed genes acting on them. Rows and columns are clustered based on hierarchical clustering in the leftmost plot. For the left plot, all effects with FDR q-value > 0.2 are whited out, whereas this threshold is relaxed to 0.5 for the middle and right plots. (**E**) (Left) Scatterplot of all significant effects (FDR q < 0.05; N = 19,909) from the cell-pooled screen (X-axis) versus the same effects in the conventional screen (Y-axis). Effects are represented in terms of log-fold changes in expression relative to control cells. R, Pearson’s correlation. SC, sign concordance. (Right) Held-out validation accuracy of down-sampled conventional screen and cell-pooled screen. X-axis: Size of down-sampled conventional screen by number of channels. Y-axis: Validation accuracy (in terms of correlation of effects with held-out validation dataset) of top 19,909 effects from the down-sampled conventional screen. Effects are estimated using FR-Perturb, elastic net regression, or negative binomial regression. The same method is used to estimate effects in both the down-sampled conventional data and validation data. The results from the cell-pooled screen (dotted line) are shown for estimation with FR-Perturb only (see **Figure S2D** for results using other methods). (**F**) (Left) Precision-recall curves computed from down-sampled conventional screen and cell-pooled screen (dotted line). True positives are the 79,100 significant effects from the held-out validation dataset, while the classification threshold being varied (X-axis) is the significance of the effects in the down-sampled conventional screen. All effects displayed are learned using FR-Perturb. (Right) AUPRCs (Y-axis) computed from the down-sampled conventional experiment when varying the number of channels (X-axis). Effects are computed using all three inference methods.

In the guide-pooled experiment, we infected cells expressing dCas9-KRAB at high MOI (**Methods**) and profiled a single cell in each droplet across seven channels, while for the corresponding conventional Perturb-seq we infected cells with the same guide library at low MOI and analyzed 19 channels. From the guide-pooled experiment, we obtained 24,192 cells after filtering (conventional: 66,283), where 35% of the cells (8,448) contained three or more guides (**Figure 4A)**, with 2.50 guides on average per cell (conventional: 1.13) (**Figure 4B**) and 101 cells containing a guide for each perturbed gene on average (conventional: 115) (**Figure 4C**). We measured expression for 16,268 total genes (conventional: 18,617). The guide-pooled screen was approximately 3 times cheaper than the conventional screen.

**Figure 4.**
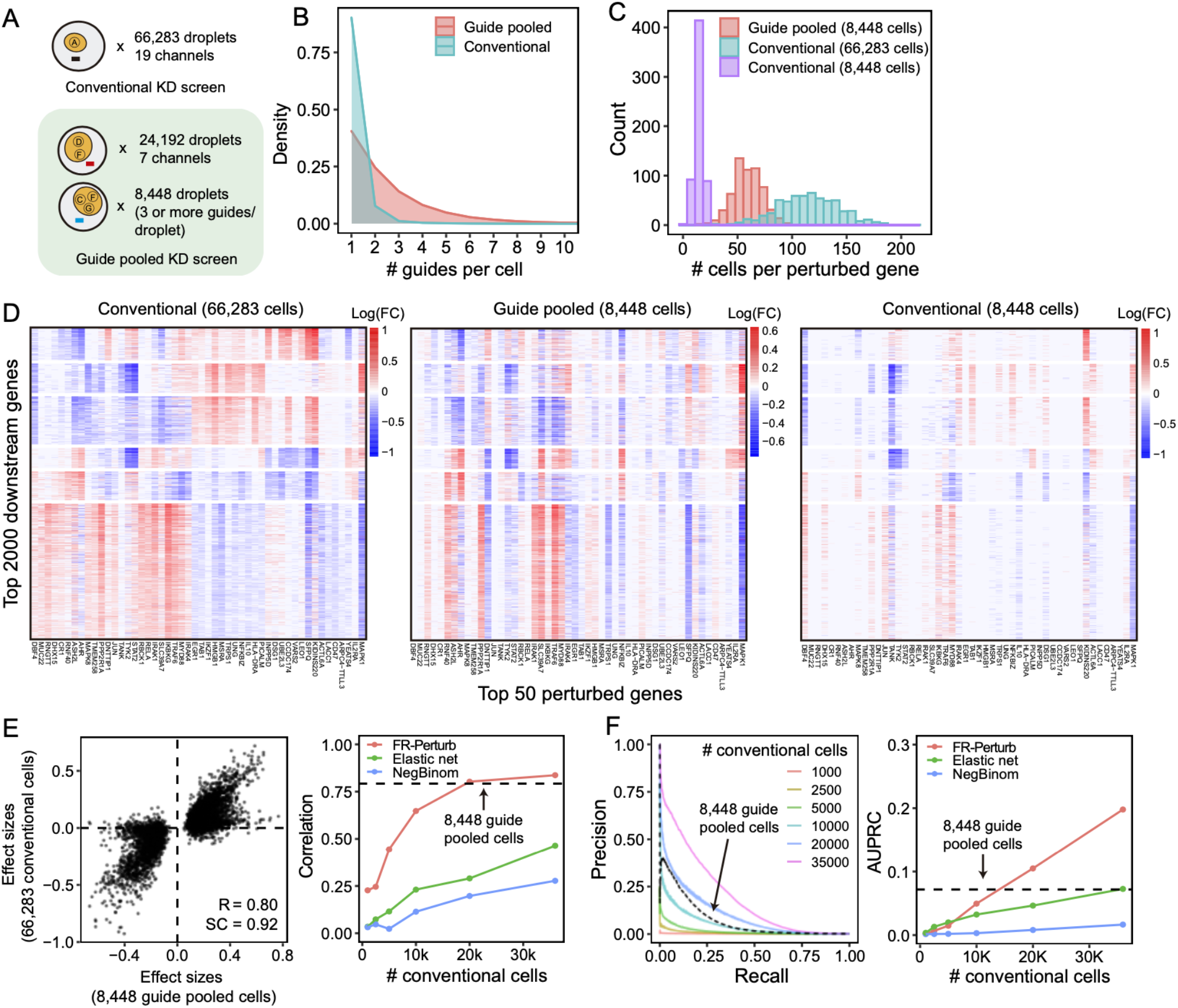
Evaluating guide-pooled Perturb-seq versus conventional Perturb-seq. (**A**) Number of channels and droplets from the conventional validation screen (top) and guide-pooled screen (bottom). We focus our analysis on the subset of 8,448 droplets from the guide-pooled screen with at least 3 guides per droplet. (**B**) Distribution of cells based on # of guides they contain for the full guide-pooled and conventional screens. In practice, we only directly measure the # of guides/droplet rather than guides/cell, but these quantities are equivalent given 1 cell/droplet. (**C-F**) See captions for **Figure 3C-F.**These analyses were conducted in an identical fashion, with the only difference that the screens are down-sampled based on cell count rather than channel count.

### Compressed Perturb-Seq by cell pooling yields accurate perturbation effects at high efficiency

The perturbation effect sizes estimated by Perturb-FR from the cell-pooled Perturb-seq screen (**Methods**) agreed well with its conventional counterpart (**Table S2**). When estimating effects, we included read count, cell cycle, and proportion of mitochondrial reads as covariates^2^, and we combined sgRNAs targeting the same gene while retaining the subset of sgRNAs for a gene with maximal concordance of effects across random subsets of the data (**Methods**). The significant effects from the compressed experiment (N = 19,909) were strongly correlated with the corresponding effects from the conventional experiment (Pearson’s R = 0.92, sign concordance = 0.96, **Figure 3E**). Notably, we observed many more significant effects overall in the conventional screen than the cell-pooled screen (216,220 *vs*. 19,909; FDR q-value < 0.05), but this is expected given that we intentionally generated a much larger and more highly powered conventional screen to enable data splitting and cross validation analyses (see below).

The cell-pooled experiment yielded substantially more signal per experimental unit (channel) than the conventional one (**Figure 3D-F**). First, the global clustering of effects learned from a single cell-pooled channel was much less noisy than from a single conventional channel (**Figure 3D**). Moreover, approximately four conventional channels were needed to obtain the same number of significant effects as one cell-pooled channel (**Figure S2A**). Next, to quantitatively assess the specificity of each approach, we held out half of the conventional data as a validation set, then we down-sampled the remaining half to different numbers of channels and compared the top 19,909 most significant effects learned from the down-sampled data (matching the number of significant effects in the cell-pooled screen) to those in the held-out validation set. We found that 5-6 conventional channels were needed to achieve equivalent validation accuracy (correlation) as one cell-pooled channel (**Figure 3E**). The relative efficiency gains of the compressed screen were consistent when varying the number of effects being compared (**Figure S2C**). We also assessed the sensitivity of each approach by testing whether the significant effects determined from the validation set were recovered by the down-sampled conventional or cell-pooled screens. We constructed precision-recall curves, calling “true positives” the 79,100 significant effects from the validation dataset and varying the classification threshold by the significance of the effects in the down-sampled conventional or cell-pooled datasets. One cell-pooled channel had comparable AUPRC to 4 conventional channels (**Figure 3F**), with consistent efficiency gains when varying the number of true positive effects (**Figure S2C**).

Moreover, FR-Perturb substantially outperformed the established inference methods we tested: elastic net regression^2^ and negative binomial regression^16^. Repeating the same analyses as above with each method (**Methods**), the concordance between the down-sampled conventional data and validation data, and between cell-pooled and conventional data, was much higher with FR-Perturb than prior methods (**Figure 3E-F, Figure S2D**). By down-sampling the cell-pooled screen, we found that ∼1/5 of a cell-pooled channel analyzed with FR-Perturb achieved the same validation accuracy as 10 conventional channels analyzed with existing methods (**Figure S2B**). We assess the cost savings of cell pooling over the conventional approach while factoring in sequencing costs in the **Discussion**.

### Guide pooling yields accurate perturbation effects at high efficiency

Guide-pooled Perturb-seq was also concordant with its conventional counterpart, based on a similar evaluation scheme as above. For the guide-pooled screen, we focused on the 8,448 cells with 3 or more guides. This number of guides per cell can be achieved with sequential transduction, as done for 2 of the 7 channels (**Methods, Figure S1B**). We learned perturbation effects from both screens using FR-Perturb, with slight modifications to account for differences in the guide-pooled *vs*. cell-pooled screens (**Methods, Table S2**). The 5,836 significant effects from the guide-pooled cells were strongly correlated with the same effects from the conventional screen (Pearson’s R = 0.80, sign concordance = 0.92) (**Figure 4E**). Thus, even if some non-linear effects exist between guides, the overall assumption of additivity holds broadly enough to infer many accurate effects. As with the cell-pooled screen, the total number of significant effects was much lower in the 8,448 guide-pooled cells *vs*. the full conventional screen (5,836 *vs*. 95,526; q-value < 0.05), but this is expected because our conventional screen was by design much larger and more highly powered overall to enable down-sampling analyses.

The guide-pooled screen was substantially more efficient than the conventional screen per experimental unit (cell), and FR-Perturb provided more accurate effect sizes than established methods. Around 2.5x more conventionally studied cells were needed to obtain the same number of significant effects as guide-pooled cells (**Figure S2E**). Globally, the effect size patterns learned from the same number of cells (8,448 cells) were much less noisy in the guide-pooled screen than in the conventional screen (**Figure 4D**). Approximately twice as many conventional cells were required to learn effect sizes at the same correlation (**Figure 4E**) or to attain the same AUPRC (**Figure 4F**) as guide-pooled cells when comparing to a held-out validation set. This relative efficiency gain was consistent when varying the number of compared effects (**Figure S2G**). Moreover, the effect sizes inferred by FR-Perturb had substantially better validation accuracy than those from the two established inference methods in both the guide-pooled and conventional data (**Figure 4E,F, Figure S2H**). We found that around 3,200 guide-pooled cells analyzed with FR-Perturb achieved the same validation accuracy as 36,000 conventional cells analyzed with existing approaches (**Figure S2F**), leading to an approximately 10-fold cell count and cost reduction over existing experimental and computational approaches (**Discussion**).

### Guide pooling is the more impactful compression approach

We next considered the strengths and limitations of the two compressed designs relative to conventional designs, and to each other.

Interestingly, cell pooling performed worse than the conventional approach on a per-droplet basis in both held-out real data and simulations (**Methods, Figure S3A**), but we still observed overall efficiency gains from cell pooling (**Figure 3D-F**) because it provided 7-fold more non-empty droplets per channel (rather than providing more cells per droplet per se). Thus, the reduced per-droplet power (which occurs due to dampening of perturbation effect sizes in a manner proportional to the number of cells in a droplet, **Supplementary Note**) is balanced by increased per-channel power from obtaining more non-empty droplets. We cannot confidently predict the degree of cell pooling needed to achieve the optimal balance between these counteracting effects. For plate-based assays such as sci-Plex^29^, our results suggest that pooling cells will decrease per-sample performance without a counteracting gain from obtaining more samples.

Conversely, the efficiency of the guide-pooled (high MOI) design readily scales with the number of guides per cell, with the best performance attained by cells with 4 or more guides in both held-out real data and simulations (**Figure S3B**). Thus, a higher number of guides per cell (achievable via sequential infections or the use of Cas12 to knock out/down multiple genes with a single construct^30,31^ should further improve efficiency, regardless of whether the assay is droplet-based or plate-based.

These observations have important implications for existing Perturb-seq screens, each of which already has some overloaded droplets (cell pooling) and multi-guide expressing cells (guide pooling) by chance or by design^1,2,13^. While these cells/droplets are often discarded, our results suggest that these cells/droplets can contain even more signal than the single-guide/single-cell containing ones and thus should be retained. To illustrate this, we used FR-Perturb to analyze a Perturb-seq knock-out screen of 1,130 genes in mouse bone marrow derived dendritic cells (BMDCs) (Geiger-Schuller et al.). In this screen, 519,535 droplets containing a single cell were obtained, of which 33% contained more than one guide by chance. By stratifying cells by the number of guides and comparing the learned effect sizes from FR-Perturb with a held-out validation subset of the data with single guide perturbations, we show that the accuracy of the effect sizes scales with the number of guides per cell and is highest in cells containing three guides (**Figure S3C**). Thus, by retaining all cells with more than one guide, the sample size of the experiment could effectively be doubled compared to the conventional approach that discards these cells (**Figure S3C**).

### Co-functional modules and co-regulated gene programs in the LPS response

We next leveraged the overall concordance of all perturbation data (conventional and compressed, knock-out (KO) and knock-down (KD)) to investigate the underlying regulatory circuitry of the LPS response. To maximize power, we merged droplets from the compressed and conventional screens together, then re-estimated all effects. There were 251,792 significant effects in the combined conventional and cell-pooled KO screen (131,161 effects in the combined conventional and guide-pooled KD), an increase of 16% (KD: 37%) over the conventional screen alone.

Overall, the KO and KD screens were concordant, with most of the significant effects (FDR q-value < 0.05) attributed to relatively few (∼5%) of the perturbations, each with widespread effects on many genes (**Figure 5A**). As expected, there were substantially more significant effects in the KO compared to the KD screen (251,792 *vs*. 131,161 effects), consistent with larger effects of KO on the target gene’s activity^32^. Effects significant in both screens (N = 26,362) were highly correlated between the screens (R = 0.92; sign consistency = 0.99; **Figure S4A-D**). The perturbations did not lead to new global cell states, such that profiles from perturbed (one or more targeting guides) and unperturbed (control guide) cells spanned the same low-dimensional space (**Figure 5C**). Thus, while many perturbations had significant and widespread effects, they did not yield radically altered phenotypic states, consistent with previous studies of this cellular response^2^.

**Figure 5.**
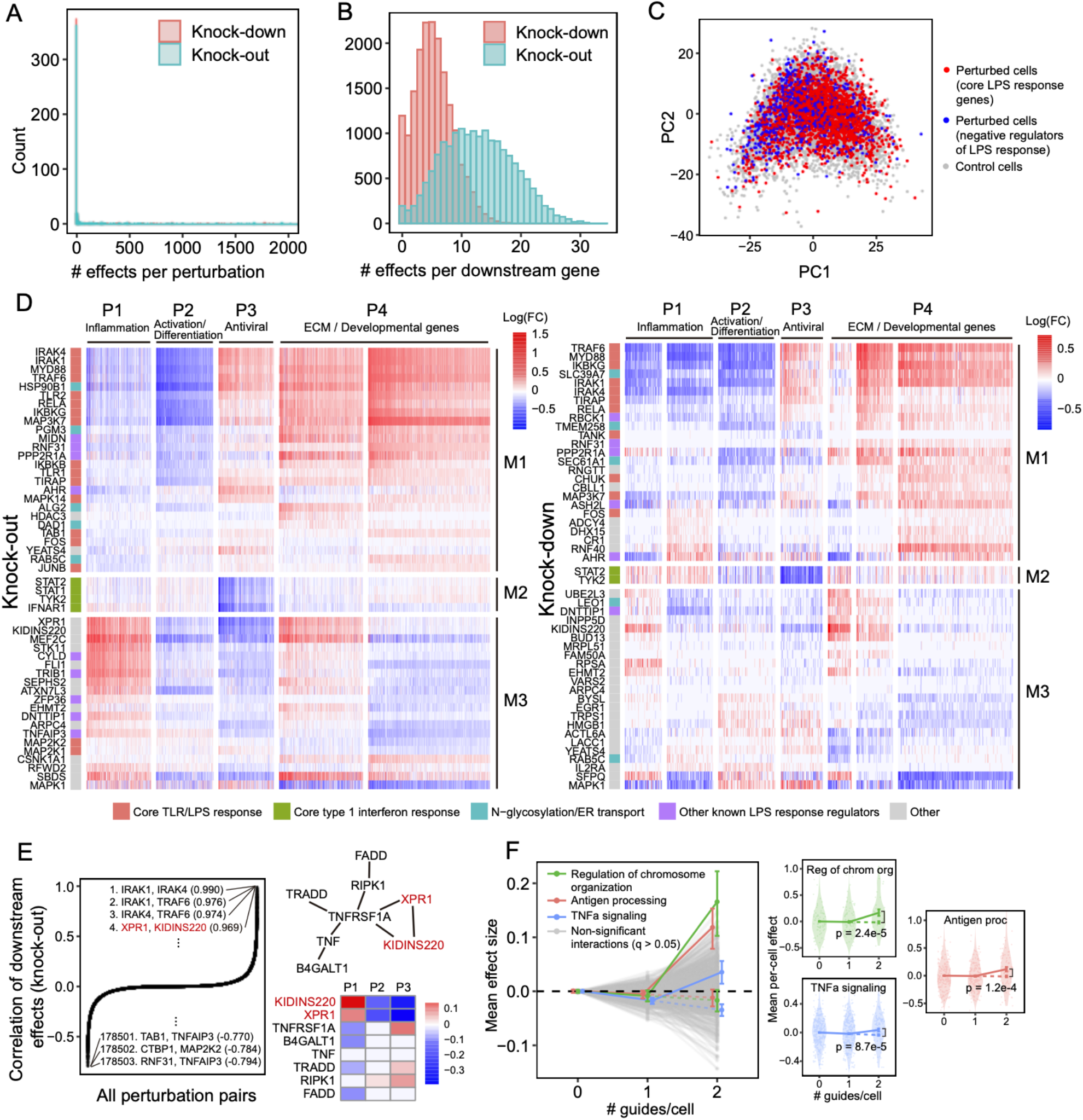
Analysis of knock-out and knock-down perturbation effects in the LPS response. (**A**) Distribution of perturbed genes based on their number of significant effects (q < 0.05) on downstream genes. (**B**) Distribution of downstream genes based on how many perturbed genes significantly affect their expression. (**C**) PCA of perturbed and unperturbed cells based on the expression of the top 2,000 most variable genes. Grey points represent cells containing a non-targeting guide only. Red points represent cells containing a guide for genes involved in the core LPS/TLR response (IKBKB, IKBKG, IRAK1, IRAK4, MAP2K1, MAP3K7, MAPK14, MYD88, RELA, TIRAP, TLR1, TLR2, and TRAF6). Blue points represent cells containing a guide for genes known to be negative regulators of the LPS response (CISH, CYLD, STAT3, TNFAIP3, TRIB1, and ZFP36). (**D**) Heatmaps of perturbation effect sizes (inferred with FR-Perturb) from the knock-out (left) and knock-down (right) screens. Rows: top 50 perturbed genes based on their average magnitude of effects on all downstream genes. Columns: top 2,000 downstream genes based on the average magnitude of effects of all perturbed genes acting on them. Rows and columns are clustered using Leiden clustering. Clusters are labelled based on their GO enrichment terms. All effects with q > 0.2 are whited out. (**E**) (Left) Correlation of knock-out effect sizes (y-axis) between all pairs of perturbed genes (x-axis). Top and bottom gene pairs are labelled. (Top right) Graph of all perturbed genes that physically interact with XPR1 and/or KIDINS220, based on AP-MS data from Bioplex 3.0^46^. Edges represent physical interaction. (Bottom right) Mean effects of perturbed genes from top right on P1-P3. (**F**) Analysis of genetic interaction effects. (Left) Effect sizes relative to control (y-axis) of cells containing 0, 1, or 2 guides (x-axis) within each perturbation module (lines connecting three dots). Modules with significant effects (q < 0.05) are highlighted in color and labeled, with the expected effect of cells containing two guides in the module represented with a dotted line. All other non-significant modules are represented in grey. Error bars represent standard errors of effect sizes obtained from bootstrapping. (Right plots) Violin plots of the mean effects of individual cells (relative to control) containing 0, 1, or 2 guides in the three perturbation modules with significant interaction effects. Dotted line represents the expected effect relative to control of cells with 2 guides. P-values are computed from permutation testing.

We organized the perturbations and genes by clustering their effect size profiles (**Methods**), observing four broad co-regulated programs of downstream genes with correlated responses across the perturbations, and three broad co-functional modules of perturbations with correlated effects on downstream genes (**Figure 5D**).

The four major co-regulated programs were present in both the KO and KD screens (**Figure 5D**), spanning key aspects of the response to LPS: inflammation (P1; cytokine, chemotaxis and LPS response genes; **Figure S4E-F**); macrophage differentiation (P2; immune cell activation, differentiation, and cell adhesion genes); antiviral response (P3; type I interferon response genes); and ECM and developmental genes (P4) (**Table S3**). Inflammation (P1) and the antiviral response (P3) are known to be regulated by LPS signaling through AP1/NF-kB and IRF3, respectively^33^, and were mostly anti-correlated in their responses to perturbation in our screen, consistent with reports that downregulation of the inflammatory response can lead to upregulation of type I interferon response^34,35^. Inflammatory signaling is known to lead to macrophage differentiation^36^, but almost all perturbations with significant effects on inflammation (P1) (in any direction) down-regulated macrophage differentiation (P2). This suggests that additional factors beyond inflammatory signaling mediate macrophage differentiation in response to LPS^37^.

Of the three major co-functional modules, KO/KD of the first module (M1) resulted in strong down-regulation of inflammation and macrophage differentiation (P1-2) and upregulation of the antiviral response and ECM/developmental genes (P3-4) (**Figure 5D**). M1 was mainly composed of core TLR/LPS response genes and genes directly up-or downstream of the pathway^33^, including MYD88, IRAK1, IRAK4, RELA, TRAF6, TIRAP, IKBKB, IKBKG, TAB1, TANK, TLR1, TLR2, MAPK14, MAP3K7, FOS, JUNB, and CHUK. Given the known function of these genes, we expect that their KO/KD will lead to down-regulation of inflammation and macrophage differentiation (P1-2), as we indeed observed. Other genes in M1 previously shown to down-regulate TNF and the inflammatory response when knocked out^25^ included two LUBAC complex proteins (RBCK1 and RNF31), genes in the OST complex (DAD1, TMEM258) and ER transport (HSP90B1, SEC61A1, ALG2), and other genes with diverse functions (MIDN, AHR, PPP2R1A, ASH2L). M1 also included two additional ER transport genes not previously implicated in immune pathways (RAB5C, PGM3), highlighting the important role of N-glycosylation and trafficking in macrophage activation^38^.

KO/KD of the second co-functional module (M2) primarily resulted in strong downregulation of the antiviral program (P3), with weak/mixed effects on other programs. M2 comprised four genes known to be core components of the type 1 interferon response^39^ – STAT1, STAT2, TYK2, and IFNAR1 – for which downregulation of the antiviral program in response to their perturbation is expected.

KO/KD of the third and final co-functional module (M3) resulted in upregulation of inflammation (P1), downregulation of macrophage differentiation and the antiviral response (P2-3), and mixed effects on ECM/development (P4). M3 included many genes with known inhibitory effects on inflammation, including ZFP36, an RNA-binding protein that destabilizes TNF mRNA^40^, enzymes CYLD and TNFAIP3 involved in deubiquitination of NF-kB pathway proteins^41,42^, pseudokinase TRIB1 and ubiquitin ligase RFWD2 which are involved in degradation of JUN^43,44^, and RELA-homolog DNTTIP1^25^. Other genes in M3 included transcription factors (MEF2C, FLI, and EGR1), chromatin modifiers (EHMT2, ATXN7L3), and kinases (CSNK1A1, STK11).

Interestingly, two of the M3 genes with particularly strong effects on all programs did not have prior immune annotations – XPR1, a retrovirus receptor involved in phosphate export, and KIDINS220, a transmembrane scaffold protein previously reported in neurons^45^. In the KO screen, this pair of genes had the fourth highest correlation of downstream effects (R = 0.97) among all 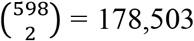 perturbation pairs (**Figure 5E**), following IRAK1/IRAK4, IRAK1/TRAF6, and IRAK4/TRAF6 which are all known to form a physical LPS signaling complex^33^. In AP-MS data^46^, XPR1 and KIDINS220 physically associate with each other and TNF receptor TNFRSF1A. Knockout of TNFRSF1A in our screen results in effects opposite to XPR1/KIDINS220 KO (**Figure 5E**), suggesting a possible inhibitory effect of this complex on TNFRSF1A.

### Guide pooling reveals second-order genetic interactions

Genetic interactions (non-additive effects) between two or more genes can in principle be inferred from cells containing two or more guides, which are generated by chance when transducing cells at low or high MOI. (**Figure 4B**). Here, guide-pooling can provide increased efficiency compared to the conventional approach, like in the first-order case (**Supplementary Note**).

We first attempted to estimate second-order interaction effects and their p-values from the guide-pooled screen and corresponding conventional KD screen by adding interaction terms to the perturbation design matrix (**Methods**). However, although we could generate point estimates of second-order effects^2^, none of these effects was significant in either screen due to insufficient power (**Figure S5A**), even with a lax significance threshold (q < 0.5).

To increase power, we aggregated perturbations into modules defined by GO annotations (**Table S4A**) and learned the overall impact of second-order interactions within and between each module on each gene program (**Methods**). Here, we define an interaction effect as the deviation from the sum of first-order effects for cells that contain any two perturbations from either the same module (intra-module interactions) or two different modules (inter-module interactions) (**Methods**). To ensure adequately sized groupings, we aggregated perturbations into 490 (possibly overlapping) modules each with at least 20 genes, such that any pair of perturbations in each module was represented in an average of 87 cells in the guide-pooled screen (conventional: 30 cells) (**Figure S5B**). We also constructed 30 non-overlapping modules by clustering the original 490 modules (**Methods**), resulting in 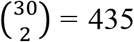 module pairs among which we could compute inter-module interactions. To increase power, we grouped downstream genes by their program (P1-4) membership (**Figure 5D**), computing mean effects on these four programs rather than on individual genes. The results from this analysis represent the extent of intra- and inter-module interactions on each key program.

We detected three co-functional modules with significant (q < 0.05) intra-module interaction effects on at least one program from the guide-pooled screen (**Figure 5F; Table S4B**), while we detected no significant interactions from the substantially larger conventional screen (even at q < 0.5) (**Figure S5C; Table S4C**). Two of the significant interaction effects – with genes for regulation of chromosome organization (p = 2.4×10^−5^) and antigen processing (p = 1.2×10^−4^) – had insignificant first-order effects on the antiviral program (P3), while having significant positive second-order effects. The third, TNFa signaling, had a significant negative first-order effect on the inflammatory/LPS program (P1) (p = 2.0×10^−4^) and significant positive second-order effect (p = 8.7×10^−5^). This effect is consistent with the reported non-linear relationship between gene dosage and TNF signaling activity when comparing heterozygous versus homozygous KO mice for either TNF^47^ or the TNF receptor TNFRSF1A^48^. Interestingly, we did not observe any significant inter-module interactions from either screen (**Figure S5D; Table S4D**,**E**), which may suggest that perturbations in different modules are less likely to interact with each other^49,50^.

### Integration of perturbation effects with genome-wide association studies implicates disease-relevant perturbations

Because dysregulation of innate immune responses plays a key role in many human diseases^51^, we next asked whether the perturbation effects learned from our *in vitro* screens can help identify disease-relevant genes and processes. *In vitro* screens may be especially helpful for this aim given that many of the perturbed genes from our screens are under strong selective constraint in human populations (**Figure S6A**), making them challenging to directly connect to disease through genome-wide association studies^52^ (GWAS) due to fewer common variants in or around the gene^53,54^. To investigate this, we obtained summary statistics from GWAS of 64 distinct human diseases and traits (**Table S5A**), including autoimmune diseases and blood traits, as well as non-immune traits/diseases (*e*.*g*. height, BMI, schizophrenia, type 2 diabetes). Using sc-linker^55^, we computed the overall heritability enrichment of these 64 traits/diseases in SNPs in/around genes comprising perturbation modules M1-3 (**Methods)**. We observed significant heritability enrichment (p < 0.001) for M3 (genes that suppress the LPS response) for two blood traits (lymphocyte and neutrophil percentage), but did not observe significant enrichment for M1 (positive regulators of the LPS response) or M2 (genes involved in the antiviral response) for any traits (**Figure S6B**).

Instead, we hypothesized that if a perturbed gene is important for disease, then disease heritability may be enriched near the *downstream genes* it affects^12,56^. To test this hypothesis, we constructed two “perturbation signatures” for each perturbed gene that include all genes that are significantly upregulated (“negative” targets) or downregulated (“positive” targets) by its KO/KD. We retained signatures with at least 100 genes, resulting in a total of 1,634 perturbation signatures from both the KO and KD screens. We also constructed signatures corresponding to the gene programs P1-4 (**Figure 5D**). As above, we used sc-linker to test for disease heritability enrichment for each signature/phenotype pair (**Methods**).

23 signatures associated with 16 perturbed genes had significant heritability enrichment scores for at least two phenotypes (p-value < 0.001). Meanwhile, 7 phenotypes that all reflect immune or blood traits (inflammatory bowel disease, eczema, rheumatoid arthritis, asthma, primary biliary cirrhosis, and eosinophil percentage) had significant scores for at least two perturbation signatures (**Figure 6A**; **Figure S6C**,**D**; **Table S5B**,**C)**. Most of the significant signatures (15/23) were from the KO screen, suggesting that the expression effects from KO are more suited for this analysis (either because they are more disease-relevant or more powered due to capturing more effects). Among the downstream programs P1-4, we observed significant enrichment from only P2 on three immune traits: inflammatory bowel disease, eczema, and primary biliary cirrhosis (**Figure S6B)**.

**Figure 6.**
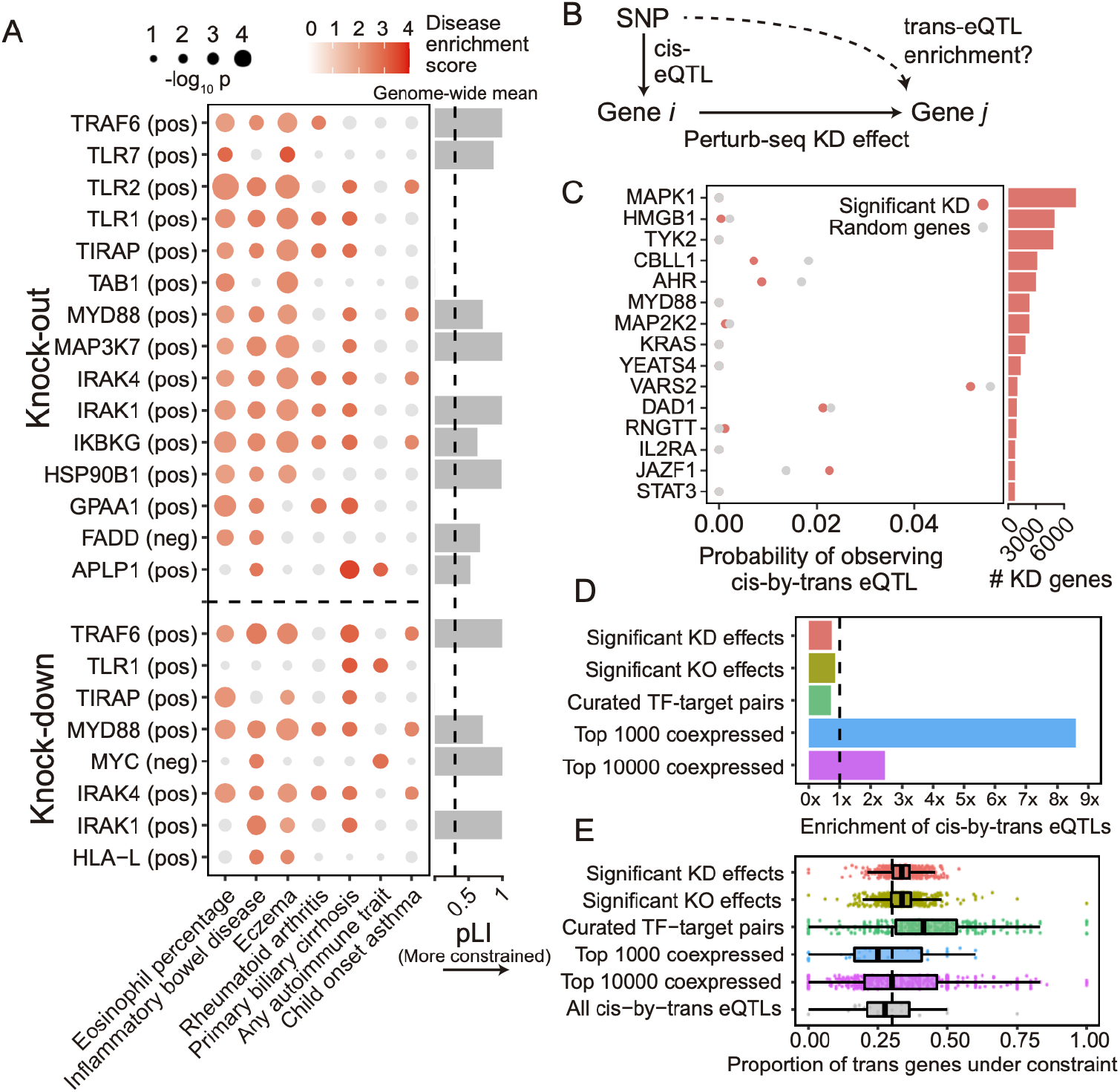
Integration of population-genetic screens with Perturb-seq. (**A**) Heritability enrichment scores of signatures comprising genes significantly modulated by perturbations (rows) across human traits (columns), computed using sc-linker^55^. “Pos” indicates the set of genes whose expression changes in the same direction as the perturbed gene (*i*.*e*., downregulated by the perturbation), with the opposite applying to “neg”. Displayed are all perturbation signatures and traits with at least 2 significant (p < 0.001) effects. Non-significant scores are greyed out. (Barplot) Probability of loss-of-function intolerance^54^ (pLI) of the corresponding perturbed gene. (**B**) Schematic of eQTL integration analysis, aiming to test whether *trans*-regulatory relationships learned from Perturb-seq are also present in eQTL studies. For all gene pairs in which gene *i* exerts an effect on gene *j* (*i*.*e*., has a significant knock-down effect in our Perturb-seq screen), we would expect that gene *i* and gene *j* are enriched for *cis*-by-*trans* eQTLs. (**C**) Using data from an eQTL study closely matching our cell type and treatment^26^, shown is the probability of observing significant *cis*-by-*trans* eQTLs among the top 15 perturbed genes from our knock-down screen and their affected downstream genes (red) compared to random downstream genes (grey). (**D**) Enrichment of significant *cis*-by-*trans* eQTLs among various sources of gene-gene pairs: significant KO/KD effects (representing significant gene-gene effects from our KO and KD screens, respectively), curated transcription factor (TF) and target gene pairs^61^, and the top 1,000/10,000 most co-expressed gene pairs (based on correlation of expression across samples) from the eQTL dataset. Enrichment is computed relative to random *trans* genes for each *cis* gene, then averaged over all *cis* genes. (**E**) Selective constraint on *trans* genes from (D) plus all significant *cis*-by-*trans* eQTLs from the Fairfax dataset. Each point represents a *cis* gene, while the x-axis represents the proportion of the *trans* genes for each *cis* gene that are under selective constraint (determined as having a pLI > 0.5).

Most of the significant signatures (17/23) were from genes in core LPS and TLR signaling pathways that fall into perturbation module M1 (even though M1 did not exhibit any direct heritability enrichment itself; **Figure S6B**): TRAF6 (positive), TLR7 (positive), TLR2 (positive), TLR1 (positive), TIRAP (positive), TAB1 (positive), MYD88 (positive), MAP3K7 (positive), IRAK4 (positive), IRAK1 (positive), and IKBKG (positive). Other significant signatures include HSP90B1 (positive), an ER transport gene important for innate immunity^57^ that is co-functional with the core LPS genes (**Figure 5D**); FADD (negative), a pro-apoptotic gene downstream of LPS signaling that serves for negative feedback^33^; MYC (negative), an oncogene with known immunosuppressive effects^58,59^; and poorly characterized pseudogene HLA-L. The two remaining significant signatures are for genes whose functions are not previously associated with the immune system, including APLP1 (an amyloid beta precursor-like gene primarily involved in brain function that interestingly contains a missense variant associated with severe influenza^60^) and GPAA1 (involved in anchoring proteins to the cell membrane). Thus, by leveraging gene-gene links learned from our screens, we were able to identify disease-relevant genes that we were underpowered to detect through direct heritability analyses (**Discussion**).

### Perturbation effects do not overlap with *trans*-regulatory links from an eQTL study in primary monocytes

*Trans*-genetic gene regulation (*i*.*e*., regulation of gene expression distal to the given SNP) has been proposed as a primary mediator of genetic effects on human disease^62^. *Trans*-genetic gene regulation can be studied through either population-level genetic data (via expression quantitative trait loci (eQTL) studies^63,64^), or through experimental perturbation of gene expression^12^, such as the screens conducted in our study. Although both types of data can in principle be used to learn the same *trans* effects, their consistency with each other has not been empirically evaluated.

We therefore compared gene-gene regulatory links between our Perturb-seq screen and a *trans*-eQTL analysis in primary patient-derived monocytes treated with LPS^26^ (N = 432), closely matching our cell line. We define a gene-gene regulatory link in eQTL studies based on *cis*-by-*trans* colocalization, where a *cis*-eQTL for gene *i* is also a *trans*-eQTL for gene *j* via a (presumed) *trans*-regulatory effect of gene *i* on gene *j* (**Figure 6B**). Here, we assume that a perturbation of a *cis*-eQTL on the expression of gene *i* is analogous to the experimental KD in our system. We used coloc^65^ to compute the posterior probability of *cis*-by-*trans* colocalization, while accounting for linkage disequilibrium between SNPs (**Methods**). To determine whether the regulatory links learned for a given perturbed gene *i* from Perturb-seq are reflected in the eQTL analysis, we compared the proportion of downstream genes *j* of gene *i* in Perturb-seq that colocalize with gene *i* in the eQTL study, *P*(*coloc* _*gene i* → *gene j*_), with the proportion of random expressed genes that colocalize with *i, P*(*coloc*_*gene i* → *gene j*_) (**Methods**).

Surprisingly, *P*(*coloc*_*gene i* → *gene j*_) was slightly lower than *P*(*coloc*_*gene i* → *gene j*_) for individual perturbed genes *i* (**Figure 6C**), as well as when aggregating across all perturbed genes (**Figure 6D**). Moreover, we observed no relationship between either the significance or magnitude of the effect of gene *i* on gene *j* and *P*(*coloc*_*gene i* → *gene j*_) (**Figure S7A**). We observed similar negative results when obtaining gene-gene links from our KO data or from a curated list of transcription factor-target gene pairs^61^ (**Figure 6D**). Using an alternative way of quantifying gene-gene links in eQTL studies that does not make assumptions about the number of causal variants (*i*.*e*., bivariate Haseman-Elston regression to estimate genetic correlation of expression^66^; **Methods**) yielded similar results (**Figure S7B**,**C**).

Conversely, we did observe significant enrichment of *cis*-by-*trans* eQTLs in gene pairs co-expressed in the same eQTL study (**Figure 6D**), as has been observed in other trans-eQTL studies^63^. Notably, co-expression in eQTL datasets is dominated by environmental effects rather than genetic effects^67^. Thus, given that the two effects are independent across samples, we would not ordinarily expect the most strongly co-expressed genes to be enriched for *cis*-by-*trans* eQTLs, suggesting that they may be confounded in part by unmodelled technical artifacts or inter-cellular heterogeneity (**Supplementary Note**). We also observed that the level of negative selection on the *trans* gene mirrored the patterns of *cis*-by-*trans* eQTL enrichment (or lack thereof) we observed in the previous analyses (**Figure 6E**), suggesting that our power to detect *cis*-by-*trans* eQTLs was affected by selection-induced depletion of SNPs affecting the *trans* genes^54,68^ (**Supplementary Note**).

## DISCUSSION

Here, we evaluated a new approach for conducting Perturb-seq based on generating composite samples, which involves either overloading microfluidics chips to generate droplets containing multiple cells (cell pooling), or infecting cells at high MOI so that each cell contains multiple guides (guide pooling). We also propose a new method, FR-Perturb, to estimate perturbation effect sizes from composite samples, which increases power by estimating sparsity-constrained effects on latent gene expression factors rather than on individual genes. We tested our approach by perturbing 598 immune-related genes in a human macrophage cell line. We found that our experimental approaches of cell pooling and guide pooling, combined with the use of FR-Perturb to infer effect sizes, lead to large cost reductions over conventional Perturb-seq while maintaining the same accuracy. Guide pooling also significantly increases power to detect genetic interaction effects and reduces the number of cells needed for screening.

Inference with FR-Perturb leads to substantially improved out-of-sample validation accuracy over conventional gene-by-gene methods (*e*.*g*., elastic net, negative binomial regression) in both conventionally generated data and compressed data. FR-Perturb is thus useful for inferring effects in any type of Perturb-seq screen, even conventional screens that do not adopt our proposed experimental changes. The improved performance of FR-Perturb in both conventional and compressed settings likely stems from perturbation effect sizes being inferred on latent gene expression factors that aggregate many co-expressed genes, thereby denoising the expression counts of individual genes which are especially noisy/sparse in single-cell data.

Compressed Perturb-seq using cell-pooling led to a 4 to 20-fold cost reduction over existing approaches in our experiments, while guide-pooling led to a 10-fold cost reduction, based on matching the number of samples needed to obtain equivalent out-of-sample validation accuracy. These numbers factor in both our experimental changes (cell-pooling or guide-pooling) combined with the use of FR-Perturb to infer effects, compared to existing experimental approaches combined with the use of existing methods to infer effects (elastic net or negative binomial regression). For cell-pooling, we account for the fact that cell-pooled channels require deeper sequencing than conventional channels (4x per channel in our case): assuming that one cell-pooled channel is equivalent to 10 conventional channels and costs are equally divided between library preparation and sequencing in the conventional screen (as was the case in our screen), this implies a relative cost of 0.5 * (0.1x library preparation costs + 4 * 0.1x sequencing costs) = 0.25x, or a 4x reduction, of cell pooling. We also provide an estimate based on our analysis that only 1/5 of a cell-pooled channel is sufficient to achieve the same validation accuracy as 10 conventional channels, resulting in a relative cost of 0.5 * (0.2 * 0.1x library preparation costs + 0.2 * 4 * 0.1x sequencing costs) = 0.05x, or a 20x reduction. Because we did not generate enough conventional data to match the performance of a single cell-pooled channel (which cannot be sub-divided in a real experiment), we report the cost reduction from cell pooling as a range rather than an exact estimate. On the other hand, guide pooling produces composite samples at the level of cells and does not require increased per-channel sequencing costs, so the cost reduction can be directly computed as the ratio of required conventional cells to guide-pooled cells (10x in our case).

Compressed Perturb-seq reduces costs due to RNA library preparation without altering the sequencing step of scRNA-seq. Thus, it can in principle be paired with approaches that increase the efficiency of sequencing via new technologies^69^ or targeted sequencing^70^, resulting in further improvements to the efficiency of Perturb-seq. Concurrent results also demonstrate the power of compressed screening with bio-chemical perturbations in high-fidelity cellular model systems (Mead et al., companion manuscript).

Cell pooling and guide pooling are complementary approaches with different strengths and limitations. Unlike cell pooling, guide pooling has the drawbacks that it requires that nonlinear interaction effects do not systematically bias phenotypes, and it potentially suffers from cellular toxicity caused by multiple viruses infecting each cell and/or multiple double stranded breaks. Meanwhile, unlike guide pooling, cell pooling has the drawbacks that it requires increased sequencing depth per channel to account for more non-empty droplets, and it loses per-droplet signal due to dilution of effect sizes (**Supplementary Note**). Due to the latter fact, cell pooling requires many more cells than guide pooling to achieve the same performance, which can be prohibitive in certain settings where cell count is limited^8,13^. Because guide pooling performs best with high guide number per cell (4 or more), whereas cell pooling does not perform well with high cell count per droplet, we posit that guide pooling (but not cell pooling) can be readily scaled up to very compressed designs (in which case the use of knock-down over knock-out and Cas12 over Cas9 may be desirable to avoid cellular toxicity), likely leading to even larger efficiency gains than we observed in our screens.

An additional key advantage of guide pooling over cell pooling is that guide pooling naturally allows for the study of higher-order interaction effects. In our study, we were underpowered (even with guide pooling) to detect second-order interaction effects between individual gene pairs. However, we detected significant intra-module interaction effects from the guide-pooled but not conventional screen, serving as a proof-of-concept that such signal can be detected in the guide-pooled screen, and may be further probed in more powered future experiments. The efficiency gains brought about from guide pooling can in theory counteract the exponential growth of gene combinations (given that various assumptions are satisfied), potentially making it the only tractable way to systematically study higher-order interaction effects (**Supplementary Note**).

By integrating data from GWAS, our screens highlighted perturbed genes with downstream genes enriched for disease heritability. Many of these perturbed genes are under strong selective constraint and would require up to millions of samples to detect in GWAS^71^. Thus, our analysis represents a potential way to circumvent the issue of negative selection removing GWAS signal from large-effect disease-relevant genes, a key challenge for biological interpretation of common-variant GWAS.

Gene-gene effects learned from our Perturb-seq screens were not enriched for *cis*-by-*trans* eQTLs in a closely matched cell type and treatment. Many possible explanations exist for this observation, including (1) insufficient power to detect *trans*-eQTLs in the eQTL dataset, (2) biological differences between our cell line and primary monocytes used in the eQTL study, (3) large differences in the magnitude of perturbation between experimental KO/KD and eQTLs, and (4) confounders in the eQTL dataset (**Supplementary Note**). Explanation (1) can in theory be addressed with larger *trans*-eQTL studies^63^, though such studies often suffer from issues with confounding/intercellular heterogeneity, as evidenced by very low reported out-of-sample replication accuracy and substantial overlap (>50%) of detected *trans*-eQTLs with variants known to influence cell type proportion^63^. Meanwhile, single-cell eQTL studies^72^ can potentially address explanation (4), though such studies suffer from low power relative to sample size (∼1,000 significant *trans*-eQTL effects detected from ∼1.2 million cells^72^ versus ∼200,000 *trans* perturbation effects detected from ∼100,000 cells in our screen). We propose that our compressed screen is a powerful tool to learn *trans*-effects on gene expression, while additional work is needed to fully reconcile the differences between population-level genetic screens and experimental perturbation screens.

## Author contributions

BC and AR conceived of the project and designed the experiments. LB, JB, BS, JF, and BC ran the experiments with input from AR. DY and BC implemented FR-Perturb and conducted computational and biological analyses with input from AG. CF and KD assisted with computational analyses. KGS and BE provided data for the mouse BMDC Perturb-seq. DY, AG, AR, and BC wrote the paper with input from all authors.

## Acknowledgements

We thank Atray Dixit for early discussions on efficient screens and Orit Rozenblatt-Rosen for discussions and help. BC was supported by the Broad Fellows program and a Merkin Institute Fellowship at the Broad Institute. DY was supported by the NSF Graduate Research Fellowship Program (Grant #1745303). AG was supported by R01 HG012133 and R01 HG006399. AR was supported by an NHGRI Center of Excellence in Genome Science grant (CEGS; RM1HG006193; AR), the Howard Hughes Medical Institute, and the Klarman Cell Observatory and Klarman Incubator at the Broad Institute. AR was a Howard Hughes Medical Institute Investigator when this study was initiated.

## Conflict of Interest statement

AR is a co-founder and equity holder of Celsius Therapeutics, an equity holder in Immunitas, and was a scientific advisory board member of ThermoFisher Scientific, Syros Pharmaceuticals, Neogene Therapeutics and Asimov until July 31, 2020. AR, BE, and KGS are employees of Genentech from August 1, 2020, March 10, 2022, and November 16, 2020, respectively. AR and KGS have equity in Roche. BC and AR are co-inventors on patents filed by the Broad Institute relating to Perturb-seq and compressed sensing methods of this paper.

## Methods

### EXPERIMENTAL PROCEDURES

#### Cell culture and stimulation

THP-1 cells (ATCC, TIB202) were cultured in RPMI medium (ATCC, 30-2001) supplemented with 10% FBS (ATCC, 30-2020) and 0.05mM 2-mercaptoethanol (Sigma Aldrich, M7522). Cells were maintained between 0.8 and 2 million cells per milliliter.

Cell lines for knockout (KO) and knockdown (KD) screens were engineered with lentiviral vectors containing Cas9 (pxpr311) and dCas9-KRAB (pxpr121), respectively. Viruses were prepared using a previously published protocol (https://portals.broadinstitute.org/gpp/public/dir/download?dirpath=protocols/production&filename=TRC%20shRNA%20sgRNA%20ORF%20Low%20Throughput%20Viral%20Production%20201506.pdf) and concentrated by centrifugation in a column with a cut size of 100kDa (MilliporeSigma UFC903096). Cells were transduced by spinfection as previously described (https://portals.broadinstitute.org/gpp/public/resources/protocols).

THP-1 cell lines were infected with sgRNA libraries (described below) at a multiplicity of infection (MOI) specific for each guide-pooled experiment. 12 hours after spinfection, cells and media were diluted 1:10 and cells were allowed to recover for 48h. Cells were selected with puromycin (2 μg/mL) for four days. The selected cells were differentiated into macrophages by stimulation in 20ng/mL phorbol 12-myristate 13-acetate (Sigma Aldrich, P8139-1mg) for 24 hours. Cells were then allowed to rest in normal culture medium for 48 hours before stimulation in medium containing 100ng/mL LPS (MilliporeSigma, L4391-1mg) for 3 hours.

#### Guide library production and validation

sgRNAs for the perturbed panel of genes (described below) were designed using the Crispr-Pick tool from the Broad Institute. Four distinct sgRNAs were designed for each perturbed gene. In addition, 500 non-targeting sgRNAs and 500 safe-targeting sgRNAs (*i*.*e*., guides targeting intergenic regions of the genome) were included. Oligonucleotide libraries were synthesized by Twist Biosciences, then amplified and inserted into a CROP-Seq vector^4^ with sgOpti scaffold (Addgene #106280) via Gibson assembly. Cloned libraries for KO, KD, and control sgRNAs (non-targeting and safe-targeting) were sequence-validated as previously described (https://portals.broadinstitute.org/gpp/public/dir/download?dirpath=protocols/production&filename=cloning_of_oligos_for_sgRNA_shRNA_nov2019.pdf). Viral libraries were produced as described above (without concentration), and an MOI was determined by transfecting cells with scaled dilutions of the virus covering a 100-fold dynamic range and quantifying survival rate after selection.

#### Conventional Perturb-Seq, cell-pooling, and guide-pooling (scRNAseq & dialout library production)

For conventional screens, the infected (MOI 0.25) and stimulated THP-1 cell suspension was prepared for droplet generation according to the manufacturer’s suggested protocol (10x Genomics, CG00053 Rev C). Channels aiming to recover 5,000-10,000 cells were loaded on the 10x Chromium Controller and the protocol was followed according to the manual for Chromium Next GEM Single Cell 3’ Reagent Kits v3.1 (CG000315 Rev C).

For cell-pooling (MOI 0.25), the standard 10x single cell 3’ RNAseq protocol (Chromium Next GEM Single Cell 3’ GEM, Library & Gel Bead Kit v3.1 PN-1000121) was run according to manufacturer’s recommendation, except the concentration of cells was increased to co-encapsulate multiple cells per droplet (250,000 cells loaded per channel).

For guide-pooling, cells were infected at an MOI of 10 before selection and stimulation, or were left to rest for 2 days after initial infection before infecting a second time at an MOI of 10 before selection and stimulation (**Figure S1B**). High MOI cells were loaded into droplets as in the conventional screens.

After the generation of double-stranded cDNA, part of the whole transcriptome amplification (WTA) product was set aside for targeted amplification to recover the perturbation barcode. 10ng of WTA from each channel were input into 8 cycles of PCR (primer 1 CTACACGACGCTCTTCCGATCT; primer 2 GTGACTGGAGTTCAGACGTGTGCTCTTCCGATCTTGTGGAAAGGACGAAACACC). The sample underwent a 1x AMPure XP Reagent SPRI clean (Beckman Coulter A63881) and was amplified for another 9 cycles with 8bp indexed PCR primers and purified with a 0.7x SPRI clean (primer 1 AATGATACGGCGACCACCGAGATCTACACTCTTTCCCTACACGACGCTC, primer 2 CAAGCAGAAGACGGCATA<u>CGAGAT</u>GTCGAGCAGTGACTGGAGTTCAGACGTGTGCT CTTCCGATCT).

### COMPUTATIONAL PROCEDURES

#### Selecting genes to be perturbed

A set of perturbed genes was compiled from several sources (**Table S1**). These included: a manually curated list of 35 canonical LPS response genes; the top 100 genes from a previous genome-wide CRISPR screen for regulation of TNF expression after LPS stimulation^25^; 100 genes identified as being a *cis* eQTL target of SNPs that were (in total) associated with *trans* eQTL effects for at least 4 downstream genes in primary monocytes treated with LPS^26^; 95 genes near high confidence variants in IBD GWAS loci^73^; 108 genes associated with Mendelian disorders identified by search for “bacterial infection” in the Online Mendelian Inheritance in Man (OMIM) database^74^ and 115 Mendelian genes similarly identified by “NF-kappa-b” search; and 173 genes reported in studies identified by a GWAS Catalog^75^ search for “infection” with diseases/traits related to liver disease and HIV-1 infection excluded.

The (perhaps surprisingly small) intersections between gene lists from these sources is depicted in **Figure S1A**. The final list of 598 perturbed genes was obtained by intersecting genes expressed in THP-1 cells with the combined list of 758 genes from all sources.

#### Generating expression and perturbation design matrix

Starting with raw Illumina BCL files from the sequencing output, the “cellranger mkfastq” command with default parameters (from the 10x CellRanger tool v6.0.1; https://support.10xgenomics.com/single-cell-gene-expression/software/downloads/latest) was used to generate FASTQ files. The “cellranger count” command with default parameters was used to align the expression reads to the GRCh38 build of the human transcriptome and generate a gene expression count matrix (see below for details on normalization of expression counts).

To generate the droplet by perturbation design matrix, paired-end reads (in FASTQ format) containing a droplet barcode and UMI on read 1 and sgRNA sequence on read 2 were aligned using Bowtie2 as follows. Read 2 reads were aligned to a reference constructed from the labeled sgRNA sequences using the --local option with default parameters, which performs local read alignment. Then, using a custom script, droplet barcodes were matched to the mapped guides for each paired-end read. A guide was called as “present” in a droplet if there were at least 5 UMIs for each droplet barcode / guide barcode pairs.

#### Inference using FR-Perturb

From the sequencing output of each of our Perturb-seq experiments, two matrices were directly generated (see above):

- *N* × *G* raw gene expression count matrix ***Y***, where *N* is the number of droplets and *G* is the number of sequenced genes.
- *N* × *P* perturbation design matrix ***X***, where *N* is the number of droplets and *P* is the total number of perturbed genes. Here, *x*_*ij*_ represents a binary indicator variable for whether droplet *i* contains a guide targeting gene *j* (we discuss below how we collapse multiple guides for the same gene). ***X*** also includes two additional columns corresponding to the presence of a non-targeting control guide and safe-targeting guide, respectively. Cells containing a non-targeting guide are treated as “control” cells (see below), while cells containing a safe-targeting guide are used to test for general effects of genome-targeting guides.

From these data, a *P* × *G* effect size matrix ***B*** is estimated, where *β*_*ij*_ represents the log fold change of the expression of gene *j* relative to control expression when gene *i* is perturbed. Two slightly different versions of FR-Perturb were formulated to learn ***B*** from ***X*** and ***Y*** generated from cell and guide pooling, respectively, as follows.

##### Version 1: Composition in expression space (for cell pooling)

This scenario arises from cell pooling. The relationship between ***B, X***, and ***Y*** in a given droplet *i* is modeled as:

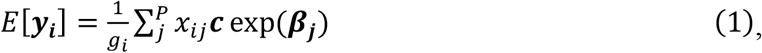

where ***y***_***i***_ is a vector of length *G* corresponding to the expression counts of all genes in droplet *i, g*_*i*_ is the number of guides contained in droplet *i* (used as a proxy for the number of cells in the droplet), *x*_*ij*_ is a binary scalar indicating whether cell *i* contains a guide for gene *j*, ***c*** is a vector of length *G* indicating the expected control expression counts of all genes, and exp(***β***_***j***_) is a vector of length *G* indicating the fold-change of expression relative to control expression for cells containing a guide for gene *j* (with ***β***_***j***_ representing the log fold-change). Note that the exp symbol here is used to distinguish fold-changes from log fold-changes, since the latter units are more commonly used to report effect sizes on gene expression. Conceptually, this model reflects the fact that expected expression measured in a droplet containing *g*_*i*_ cells is the average of the expected expression counts of the individual cells in the droplet (where the latter quantity can be expressed as ***c*** exp(***β***_***j***_) for cells containing guide *j*).

In practice, it is advantageous to model the measured expression in each droplet as the *geometric* rather than arithmetic mean of expression of the constituent cells. Simulations with real cells show that the arithmetic versus geometric mean of expression across multiple cells are very similar (**Figure S8A**), but modeling expression counts in a droplet as the latter enables us to perform inference in the space of log fold-changes rather than fold-changes. The former is symmetric around zero (whereas the latter is not) and thus leads to balanced inference of up-versus down-regulation.

Thus, Equation (1) is rewritten as follows:

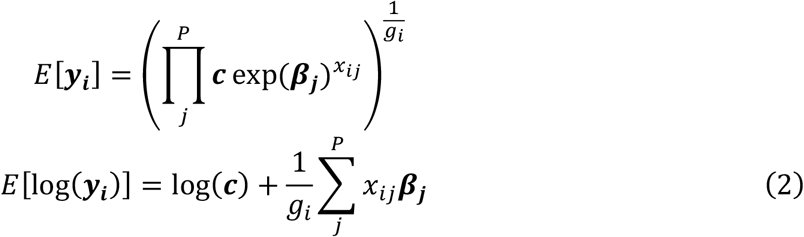

Equation (2) can be expressed simply in matrix form as *E*[***Y***′] = ***X***′***B***, where each row of 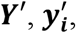, equals log(***y***_***i***_) − log(***c***), and ***X***′ is ***X*** with rows normalized to sum 1. In order to infer ***B, Y*** is transformed into ***Y***′ by taking the log(TP10K + 1) of all gene expression counts and subtracting log(***c***) from each row of *Y* (where log(***c***) represents the average log(TP10K + 1) of all genes in cells containing only non-targeting control guides). A pseudocount of 1 is included because the sparse nature of gene expression counts prevents directly taking their logarithm.

Next, the factorize-recover algorithm is applied to ***Y***′ and ***X***′ to infer ***B***. In the first “factorize” step of factorize-recover, sparse factorization is applied to ***Y***′ alone using sparse PCA, which produces *N* × *R* left factor matrix ***Ũ*** and *R* × *G* right factor matrix ***W***. *R* is a hyperparameter that controls the rank of ***Y***′. In the second “recover” step, sparse recovery is used to learn *P* × *R* matrix ***U*** from the following regression model: ***Ũ*** = ***X***′***U***, using LASSO applied to each column of ***Ũ*** (so that one column of ***U*** is learned at a time). By multiplying ***U*** by ***W*** obtained from the factorize step, a *P* × *G* matrix 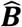 is obtained, which is an estimate of ***B***.

In practice, the magnitude of elements of 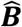 was strongly correlated with overall expression level of the downstream gene in control cells. This correlation changed (but was not removed) when varying the arbitrary pseudocount of 1 and/or scale factor of 10,000, suggesting that it was an artifact arising from log-transforming lowly expressed gene expression counts^76^. Indeed, simulations show that the magnitude of effects estimated with FR-Perturb had a negative bias that scaled with the expression level of the downstream gene, with the largest biases observed for the most lowly-expressed genes (**Figure S8C**).

This bias was removed with the following heuristic correction. First, LOESS was used to fit a curve to the plot of effect size magnitude *vs*. expression level in control cells for all entries of 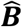. Next, all effect sizes were scaled based on the ratio of their fitted effect size magnitude from LOESS and the fitted effect size magnitude of genes with the highest expression counts (log(average TP10K) > 2). This procedure removes the global relationship between effect size magnitude and expression level of the downstream gene, while preserving heterogeneity in the average magnitude of effect sizes on individual downstream genes. In simulations, this procedure produced much less biased effect size estimates than when not scaling (**Figure S8B**,**C**).

##### Version 2: Composition in log fold change effect size space (for guide pooling)

For guide pooling data, the relationship between ***B, X***, and ***Y*** in a given droplet *i* is modeled as:

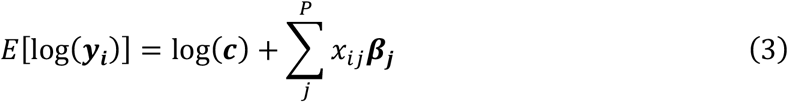

The only difference between Equation (2) and Equation (3) is the absence of the normalizing factor 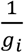 in front of the second term of the right side of Equation (3). Inference to learn ***B*** is performed as in Version 1, with the only difference that the rows of ***X*** are not normalized to have a sum of 1.

#### Covariates

Covariates corresponding to the proportion of mitochondrial reads, the total read count per cell, and cell cycle state (as determined by the CellCycleScoring function from the Seurat R package^77^) were accounted for when estimating effect sizes using FR-Perturb, by regressing the covariates out of the expression matrix according to the linear model ***Y***′ = ***CD***. Here, ***Y***′ represents the *N* × *G* normalized expression matrix (where *N* is the number of cells and *G* is the number of sequenced genes), ***C*** represents the *N* × (*C* + 1) covariate matrix including an intercept term (where *C* represents number of covariates with all covariates centered to mean 0), and ***D*** represents the fitted (*C* + 1) × *G* matrix of covariate effects on gene expression. All downstream inference was performed on the residual matrix ***Y***_*resid*_ = ***Y***′ − ***CD***.

#### Hyperparameters for FR-Perturb

The spams R package^78^ was used to perform the steps of factorize-recover, including sparse PCA and LASSO. Three hyperparameters are set in FR-Perturb: the rank *R* of ***Y***′, a tuning parameter *λ*_1_ for sparse PCA during the factorize step (which is the solution of 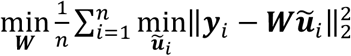 so that ∥ **ũ**_*i*_∥_1_ ≤ *λ*_1_), and a tuning parameter *λ*_2_ for LASSO during the recover step (which is the solution of min 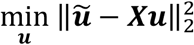 so that ∥***u***∥_1_ ≤ *λ*_2_). These were set based on maximizing cross-validation *r*^2^ as *R* = 10, *λ*_1_ = 0.1, and *λ*_2_ = 10. Analysis results were not especially sensitive to different values of *R, λ*_1_, and *λ*_2_ (**Figure S8D-F**).

#### Permutation testing for significance

Permutation testing was used to obtain two-tailed p-values for elements of 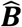. To generate an empirical null distribution for each element of 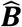, samples were permuted (*i*.*e*., rows of ***X***) and 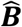 was re-inferred using FR-Perturb for each permutation. Permuting rows of ***X*** has no impact on the factorize step, since this step does not involve ***X*** (and the alternative approach of permuting rows of ***Y*** does not affect the individual factors). Thus, only the recover step was performed and ***U*** was estimated for each permutation, followed by multiplying the null ***U*** by ***W*** obtained from the factorize step to obtain the null 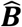 estimate. In addition, to reduce computational cost, only 500 permutations total were performed. For entries of 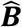 that had p-value = 0 based on these 500 permutations, a skew-t distribution was fit to the empirical null distribution for each entry using the *selm* function from the sn R package, and p-values were then re-computed for these entries from the fitted distribution. False discovery q-values were computed using the Benjamini-Hochberg procedure applied to the p-values for all entries of 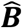.

#### Inference using negative binomial regression

Using the glmGamPoi R package^79^, ***B*** was inferred by separately running differential expression analysis for each perturbation (*i*.*e*., column of ***X***), where the two groups being compared were droplets containing only non-targeting control guides and droplets containing a guide for the perturbed gene of interest. For droplets containing multiple guides, other guides present in the droplet were ignored when forming these groups. Analytic p-values and false discovery q-values were obtained for all effect sizes from the method output.

#### Inference using elastic net

Using the spams R package^78^, the same elastic net inference procedure proposed in Dixit et al.^2^ was used to infer ***B*** from the following models: ***Y***′ = ***X***′***B*** for version 1, and ***Y***′ = ***XB*** for version 2 from above with *λ*_1_ = 0.00025 and *λ*_2_ = 0.00025 (where elastic net finds the solution to 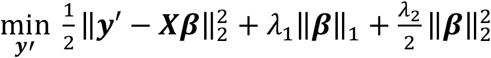 for each column of ***Y***′), matching the values used in Dixit et al. Other values for the parameters yielded similar results (**Figure S8G**). P-values for all effect sizes were obtained by permuting the rows of ***X*** a total of 10 times and re-estimating ***B*** to generate a null distribution across all values of ***B***, matching the procedure used in Dixit et al.

#### Selecting optimal guide combination for each gene

Four distinct sgRNAs were generated for each perturbed gene. When inferring effect sizes, guides were aggregated by perturbed gene to increase sample size and simplify downstream analyses. When generating the perturbation design matrix ***X***, a cell containing any guide for the gene was labelled as receiving a perturbation for the gene. However, sgRNAs have varying efficiency at KO or KD their target gene, and including guides that do not work will add noise to the effect size inference. To retain only sgRNAs that had measurable effects on their target gene, we retained guides with concordant effect size estimates across random sample-wise splits of the data (*i*.*e*., the subset of guides to the same gene showing maximal concordance).

Specifically, let *i* represent the index of a given perturbed gene, so that ***x***_***i***_ corresponds to the column of ***X*** that indicates which cells received perturbation *i*, and ***β***_***i***_ corresponds to the column of ***B*** that indicates the effects sizes on all genes’ expression from perturbing gene *i*. For each *i*, 15 different version of ***x***_***i***_ were generated, corresponding to all possible subsets of the 4 guides. For each version, any cell receiving a guide within the given subset of guides is labelled as containing a perturbation for the gene, while the remaining guides are ignored. Only ***x***_***i***_ in ***X*** was modified and the remaining columns were kept the same. Next, the dataset of interest was randomly split in half by samples (cells). FR-Perturb was used to infer effect sizes for all perturbed genes within each half. Then, the *R*^2^ of 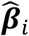 was computed between the two halves (restricting to only effects with an FDR q-value < 0.2), and the specific guide subset that produced that highest *R*^2^ was retained. The same procedure was repeated for each *i* to learn the optimal guide combination for each perturbed gene.

#### Simulations

Perturb-seq datasets were simulated at various levels of overloading using real expression counts and perturbation effect sizes estimated from our data.

##### Simulating cell-pooled data

To simulate expression data for *n* droplets containing *m* cells each, the expression of *n* · *m* cells (each containing 1 guide) were first simulated by randomly sampling control cells from our experiment, and scaling their expression counts by the fold change effect sizes of a given perturbed gene (estimated from our conventional knock-out Perturb-seq screen). A 10% probability of receiving a control guide (*i*.*e*., no change in expression) was simulated to match the proportion of control guides in the real data. Next, the expression counts of *m* cells were randomly averaged at a time to create cell-pooled data.

##### Simulating guide-pooled data

To simulate expression data for *n* cells containing *m* guides each, *m* perturbed genes were randomly selected for each cell, and the expression of a randomly selected control cell was then scaled by the product of the fold change effect sizes of the *m* perturbed genes. As before, a 10% probability of receiving a control guide was simulated.

#### Clustering and dimensionality reduction

For **Figure 5C**, dimensionality reduction was performed using PCA on the log(TP10K + 1) expression counts of all cells, where the expression values of each gene are scaled and centered to mean 0 and variance 1.

The rows and columns of **Figure 5D** were clustered using Leiden clustering^80^. First, the Euclidian distance between all pairs of genes was calculated by their perturbation effect sizes, and the FindNeighbors function from the Seurat R package^77^ was used to compute a shared nearest neighbor graph from these distances (*k*=20), followed by the FindClusters function to perform Leiden clustering on the graph with resolution parameter = 0.5, selected by visual inspection of the resulting clusters. GO enrichment analysis of the genes in the resulting clusters was performed with the ClusterProfiler package^81^ with gene sets obtained from the C2 (curated gene sets) and C5 (ontology gene sets) collections of the MSigDB^82^.

#### Learning second-order effects for individual perturbation pairs

Second-order interaction effects on gene expression in cell *i* with multiple guides were modeled as:

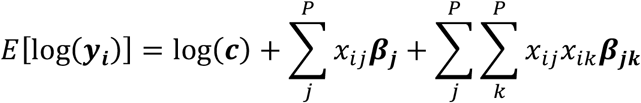

Here, log(***y***_***i***_) is a vector of length *G* corresponding to the log expression counts of all genes in droplet *i, x*_*ij*_ and *x*_*ik*_ are binary scalars indicating whether cell *i* contains a guide for gene *j* and/or gene *k*, ***c*** is a vector of length *G* indicating the expected control expression counts of all genes, ***β***_***j***_ is a vector of length *G* indicating the *first-order* effect size of guide *j* on the expression of *G* genes, and ***β***_***jk***_ is a vector of length *G* indicating the *second-order* effect size of guides *j* and *k* on the expression of *G* genes. In matrix form, the above can be represented as:

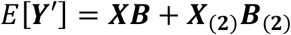

where each row of ***Y***′ equals log(***y***_***i***_) − log(***c***), ***X***_(**2**)_ is an 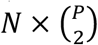 indicator matrix for whether each cell contains any of 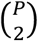 perturbation pairs, and ***B***_(**2**)_ is a 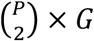 matrix of second-order interaction effects. ***B*** is known from estimating first-order effects previously, which enables the following equation to be written:

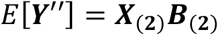

where ***Y***^′′^ = ***Y***^′^ − ***XB***. Finally, ***B***_(**2**)_ is estimated using FR-Perturb in the exact same manner as ***B***. To reduce the large size of 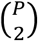, only perturbation pairs that were present in a minimum of 5 cells were included.

When estimating the significance of entries of ***B***_(**2**)_, the uncertainty in both ***B*** and ***B***_(**2**)_ must be accounted for, since the latter depends on the former. Thus, when generating a null distribution for the entries of ***B***_(**2**)_, the rows of both ***X*** and ***X***_(**2**)_ were permuted and ***B*** was re-estimated for each permutation.

#### Learning second-order effects for perturbation modules

##### Intra-modular interactions

A second-order intra-modular interaction effect was estimated for each co-functional perturbation module *M* (*i*.*e*., group of perturbed genes) on each co-regulated gene program *P* (*i*.*e*., group of downstream genes) as follows. For each pair of *M* and *P*, cells were partitioned into three sets:

1. *Control set*. Cells containing only non-targeting control guides or guides for genes without significant effects on *P*. The latter group of guides is included to increase sample size, and all these guides are collectively referred to as “control guides”.
2. *First-order set*. Cells with exactly one guide in *M*, with remaining guides in the cells falling into the “control guide” set.
3. *Second*-*order set*. Cells with exactly two guides in *M*, with remaining guides in the cells falling into the “control guide” set.

A mean expression value for *P* was computed for each set (*μ*_0_, *μ*_1_, and *μ*_1,1_ respectively) as the average standardized log(TP10K + 1) expression of all genes in *P* among the cells in the set, with covariates corresponding to read count per cell, percent mitochondrial reads, cell cycle state, and number of guides per cell regressed out of the log(TP10K + 1) expression matrix, and expression standardized to mean 0 and variance 1. The effect size of the first-order set was computed as *β*_1_ = *μ*_1_ − *μ*_0_ and the interaction effect size of the second-order set as *β*_1,1_ = *μ*_1,1_ − 2*β*_1_ − *μ*_0_. P-values for all interaction effects were computed by permuting the set membership labels of all the cells and recomputing *μ*_0_, *β*_1_, and *β*_1,1_ for the permuted sets. Standard errors for all interaction effects were computed via bootstrapping, by resampling cells from each of the sets without changing their labels.

##### Inter-modular interactions

Inter-modular interaction effects were computed using a similar approach as above. The 490 total modules were first reduced into 30 disjoint modules using Leiden clustering of a shared nearest neighbor graph defined based on the number of genes shared between gene sets. For two co-functional modules *M*_1_ and *M*_2_, the first order effects *β*_1_ and *β*_2_ were computed in the same manner as above. The second-order set was defined as cells with *at least one* guide from each of *M*_1_ and *M*_2_, with the remaining guides in the cell falling into the “control guide” category, as defined above. The mean expression of the second-order group is *μ*_1,2_. The interaction effect is defined as *β*_1,2_ = *μ*_1,2_ − *β*_1_ − *β*_2_ − *μ*_0_ and p-values and standard errors were estimated using permutation testing and bootstrapping, respectively.

#### Heritability analyses

Sc-linker^55^ was used as previously described to compute a disease heritability enrichment score for each gene set constructed from the KO and KD perturbation effect sizes or perturbation modules and gene programs. Using sc-linker, SNPs were first linked to genes using a combination of histone marks from the Epigenomics Roadmap^83^ and the activity-by-contact strategy^84^, then an enrichment score was computed for the SNPs based on the heritability enrichment of the SNPs obtained from stratified LD score regression (S-LDSC^85,86^).

More specifically, for each gene set *G*, a set of weights *A*_*G*_ = (*a*_*G*,1_, *a*_*G*,2_, …, *a*_*G,j*_) between 0 and 1 was constructed for each SNP based on the confidence of them influencing any gene in *G*, following the procedure described in Jagadeesh et al.^55^ using activity-by-contact scores^87^ and the Epigenomics Roadmap histone marks^83^ for whole blood samples. For gene sets defined from membership in perturbation modules (M1-3) or gene programs (P1-4) (**Table S3**), modules/programs were merged between the KO and KD screens. For gene sets defined based on perturbation effects, each gene was weighted by the effect size of the perturbation on the gene, normalized to lie between 0 and 1. A set of weights *A*_*all*_ = {*a*_*all*,1_, *a*_*all*,2_, …, *a*_*all,j*_} was also constructed, representing the confidence of the SNP influencing any gene across the genome. Next, heritability enrichment estimates 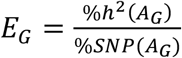 and 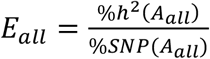 were computed for each *A*_*G*_ and *A*_*all*_, respectively, using S-LDSC. Here, 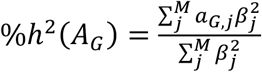 (where 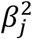 represents the squared effect size of SNP *j* on the phenotype and *M* represents the total number of SNPs), and 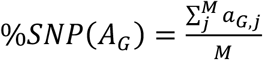. Conceptually, %*h*^2^ *G* represents the fraction of the total genetic effect on the phenotype attributed to SNPs in *A*_*G*_, while %*SNP*(*G*) represents the effective fraction of SNPs that are contained in *A*_*G*_. Thus, the ratio 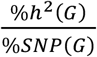 is essentially the average effect size magnitude on the phenotype for SNPs in *A*_*G*_. Finally, the enrichment score for *A*_*G*_ was computed as *E*_*G*_ − *E*_*all*_. Subtracting *E*_*all*_ controls for the baseline level of heritability enrichment for SNPs that influence any gene (since most SNPs do not influence any genes). P-values were obtained for the null hypothesis *E*_*G*_ − *E*_*all*_ = 0 using a block jackknife procedure^85^.

#### eQTL analyses

Raw genetic data for 432 European individuals and gene expression data for primary monocytes from these individuals profiled 2 hours after treatment with LPS was obtained from Fairfax et al.^26^. For each *cis*-*trans* gene pair, plink^88^ was used to compute marginal association statistics of all SNPs within 1 megabase of the promoter of the *cis* gene with the expression of both the *cis* gene and *trans* gene. All our analyses were restricted to *cis genes* with at least one significant *cis*-eQTL (q < 0.05) in the Fairfax dataset. Next, coloc^65^ was applied to the association statistics to estimate the posterior probability (with the default prior) that the *cis* and *trans* gene have a shared eQTL within 1 megabase of the *cis* gene, setting a posterior probability threshold of 0.75 to determine significant colocalization (varying this threshold does not change downstream results, **Figure S7D**). The posterior probability that each *cis* gene colocalizes with random *trans* genes was also computed. For all analyses, the top 20 principal components of the gene expression matrix were included as covariates, matching the covariates included by Fairfax et al. in their *trans*-eQTLs analysis and selected based on the fact that they maximize the number of significant *trans*-eQTLs in Fairfax et al. By restricting the *cis* gene to having a significant eQTL and comparing our effects to random genes while keeping the *cis* gene the same, we control for differences in power for detecting *cis*-by-*trans* eQTLs that arise from differential levels of selective constraint on the *cis* gene. In particular, the *cis* genes selected to be perturbed in our screens include many genes under selective constraint (**Figure S6A**), for which we have decreased power to detect *cis*-by-*trans* eQTLs compared to random *cis* genes.

Bivariate Haseman-Elston regression as implemented in the GCTA software tool^66^ was also used to compute the genetic correlation between the expression of the *cis* gene and the *trans* gene when restricting to the region 1 megabase around the promoter of the *cis* gene. Again, the top 20 principal components of the gene expression matrix were included as covariates. The method outputs a genetic correlation estimate 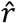 and standard error estimate 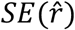 for each *cis*-*trans* gene pair. In order to obtain a combined genetic correlation estimate for all downstream genes of a given perturbed gene, all 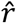 estimates were first squared and then combined using inverse variance weighing. The variance of 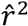 was estimated from 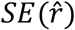 using the Delta method: 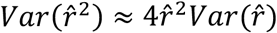.

#### Data and Software Availability

Raw and processed data for all screens were deposited in NCBI’s Gene Expression Omnibus under accession number GSE221321. Software implementing FR-Perturb can be found at https://github.com/douglasyao/FR-Perturb.

