## Supplementary Figures and Note for "Compressed Perturb-seq: highly efficient screens for regulatory circuits using random composite perturbations"

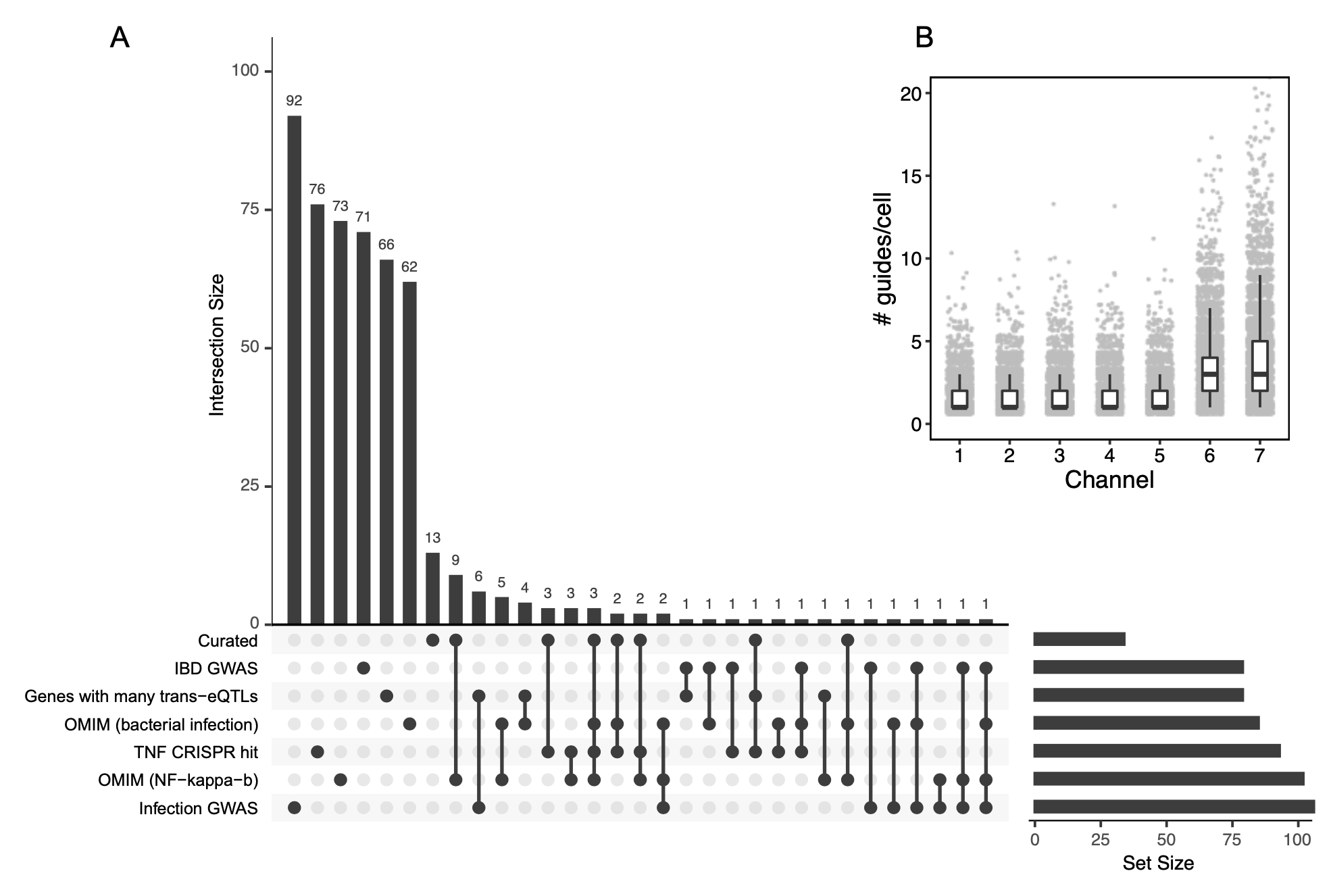

**Figure S1. Experimental design.** (**A**) Upset plot of sources of 598 genes chosen to be perturbed. Curated: manually curated list of canonical LPS response genes. IBD GWAS: genes near fine-mapped variants from GWAS of inflammatory bowel disease (Huang 2017 Nature). Trans eQTL regulator: genes with a cis-eQTL that is also a trans-eQTL for 4 or more genes in an immune model system (Fairfax 2014 Nature). OMIM (bacterial infection): genes involved in Mendelian diseases associated with the term “bacterial infection” in the Online Mendelian Inheritance in Man (OMIM) database (Amberger 2019 Nucleic Acid Research). TNF CRISPR hit: top 100 hits from a genome-wide CRISPR screen for regulation of TNF expression after LPS stimulation (Parnas 2015 Cell). OMIM (NF-kappa-b): genes involved in Mendelian diseases associated with the term “NF-kappa-b” in the OMIM database. Infection GWAS: genes associated with the term “infection” in the GWAS Catalog (Buniello 2019 Nucleic Acid Research). (**B**) Distribution of guides/cell across seven 10X channels from guide-pooled screen. Each point represents a cell, with boxes depicting the first quartile, median, and third quartile of values. For channels 6 and 7, a sequential transduction was performed, resulting in a larger number of guides per cell.

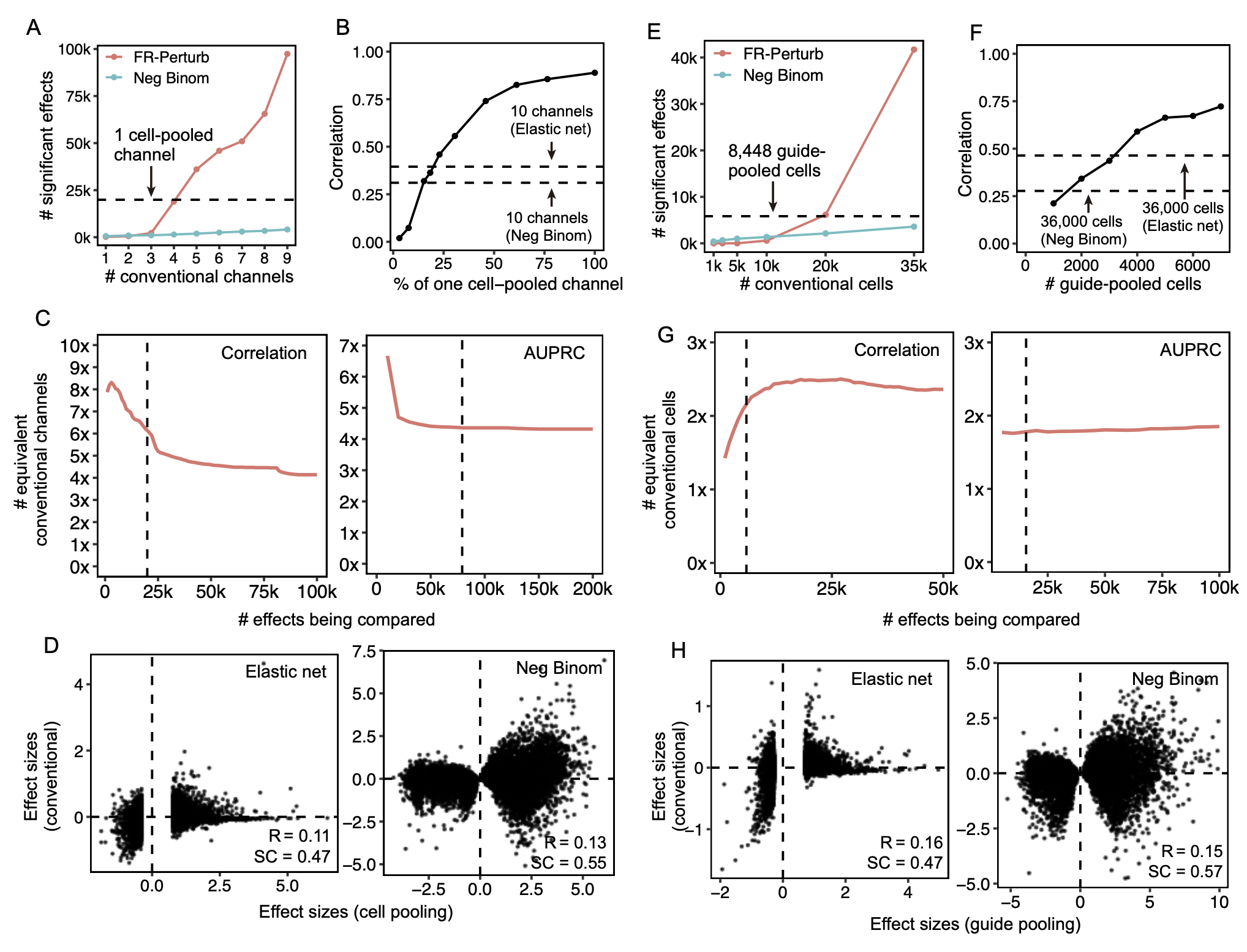

**Figure S2. Additional analyses comparing compressed versus conventional screens.** (**A**) Number of significant effects (q < 0.05) detected by FR-Perturb and negative binomial regression (y-axis) as a function of number of channels (x-axis) from the conventional knock-out screen. We do not include the number of significant effects from elastic net due to its extremely large magnitude (>1,000,000), which is inconsistent with the performance of elastic net in held-out validation analyses. (**B**) Sample size in terms of percentage of a single cell-pooled channel by droplet count (x-axis) versus out-of-sample validation accuracy (y-axis). Validation accuracy of 10 channels analyzed with elastic net or negative binomial regression is indicated with dotted lines. (**C**) Performance of cell-pooled versus conventional screen (y-axis) while varying the number of effects being compared (x-axis). Performance is quantified as the number of conventional channels needed to obtain the same correlation (left) or AUPRC (right) as one cell-pooled channel. Dotted line represents the cutoffs used in **Figure 3E-F**. (**D**) Scatterplots of top 19,909 estimated effects from the cell-pooled screen (x-axis) versus the same effects in the conventional screen (y-axis) when estimating effects using elastic net regression (left) or negative binomial regression (right). R = Pearson’s correlation, SC = sign concordance. (**E-H**) Same as **A-D**, but showing results from the guide-pooled screen (restricting to cells with 3 or more guides) and corresponding conventional screen.

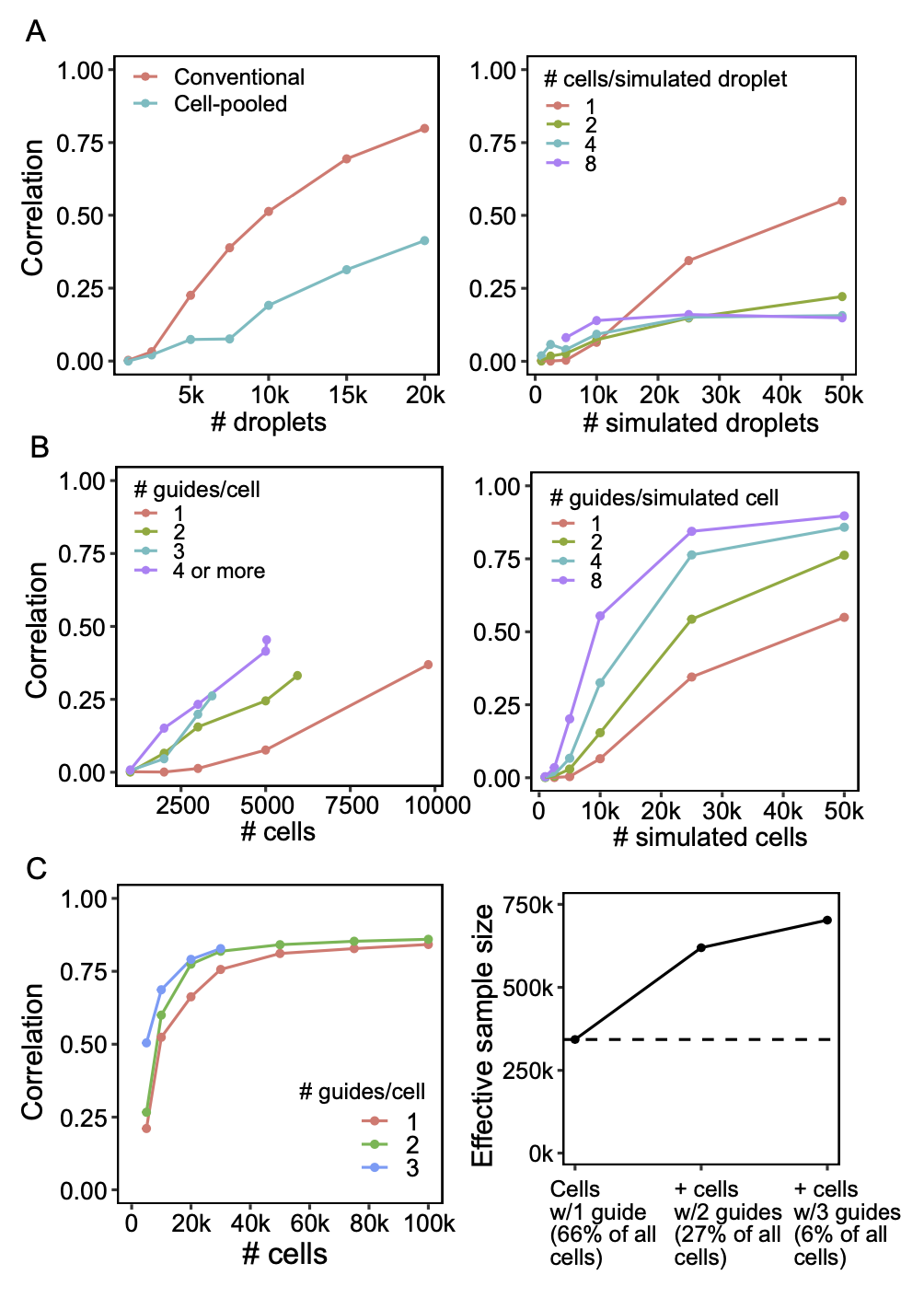

**Figure S3. Evaluating potential of compressed designs in current and future screens.** (**A**) Down-sampling droplets from cell-pooled and conventional screens. (Left) Correlation of top 10,000 estimated effects with held-out validation data (y-axis) when varying droplet count (x-axis). (Right) Correlation of top 10,000 estimated effects with true effects in simulations of cell-pooled data with varying numbers of cells/droplet. (**B**) Down-sampling cells from guide-pooled screen stratified by # guides/cell. (Left) Correlation of top 10,000 estimated effects with held-out validation data (y-axis) when varying cell count (x-axis). (Right) Correlation of top 10,000 estimated effects with true effects in simulations of guide-pooled data. (**C**) Additional signal in cells containing multiple guides (which would typically be discarded) from a conventional Perturb-seq screen of 1,130 genes in mouse BMDCs (Geiger-Schuller et al., companion manuscript). (Left) Correlation of top 10,000 estimated effects with held-out validation (y-axis) when varying cell count (x-axis). (Right) Increase in effective sample size in cells (y-axis) when including cells containing 2 or 3 guides (x-axis). Effective sample size for cells with 2 or 3 guides is computed as the number of single-guide containing cells needed to achieve the same held-out validation accuracy (from left panel).

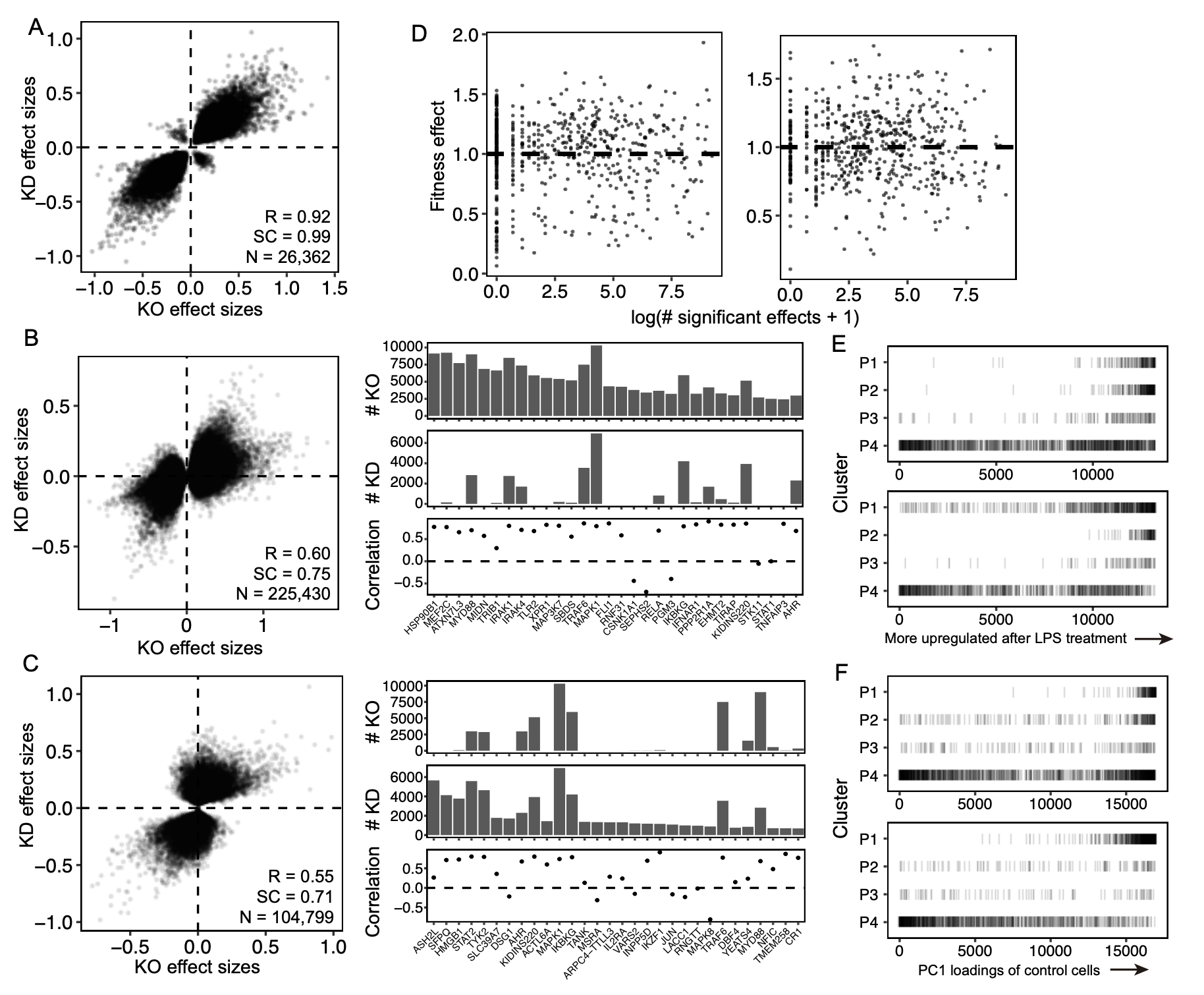

**Figure S4. Comparison of knock-out versus knock-down screens.** Scatterplots of knock-out effect sizes (x-axis) versus knock-down effect sizes (y-axis) while restricting to effects significant (q < 0.05) in both screens (**A**), effects significant in only the knock-out screen (**B**), and effects significant in only the knock-down screen (**C**). For **B** and **C**, the correlation of effects stratified by individual perturbed genes is also shown (right plots), with the number of total significant effects for each perturbed gene in the knock-out (top barplot) and knock-down (bottom barplot) screens. (**D**) Relationship between fitness (y-axis) and the number of significant downstream effects (x-axis) for each perturbed gene in the knock-out (left) and knock-down (right) screens. The fitness effect of each perturbed gene is computed as the ratio of reads between the input guide pool and the guides after infecting cells and treating with LPS (while normalizing to the guide distribution in the input pool). (**E**) Enrichment of genes upregulated after LPS treatment in the four main gene programs (**Figure 5D**) from the knock-out (top) and knock-down (bottom) screens. X-axis: genes sorted based on their expression fold changes before vs. after treatment with LPS, with more upregulated genes toward the right. Genes in the indicated gene programs are represented with a black bar. (**F**) Same as **E**, but sorting genes based on the PC1 loadings of genes from PCA performed on only unperturbed control cells.

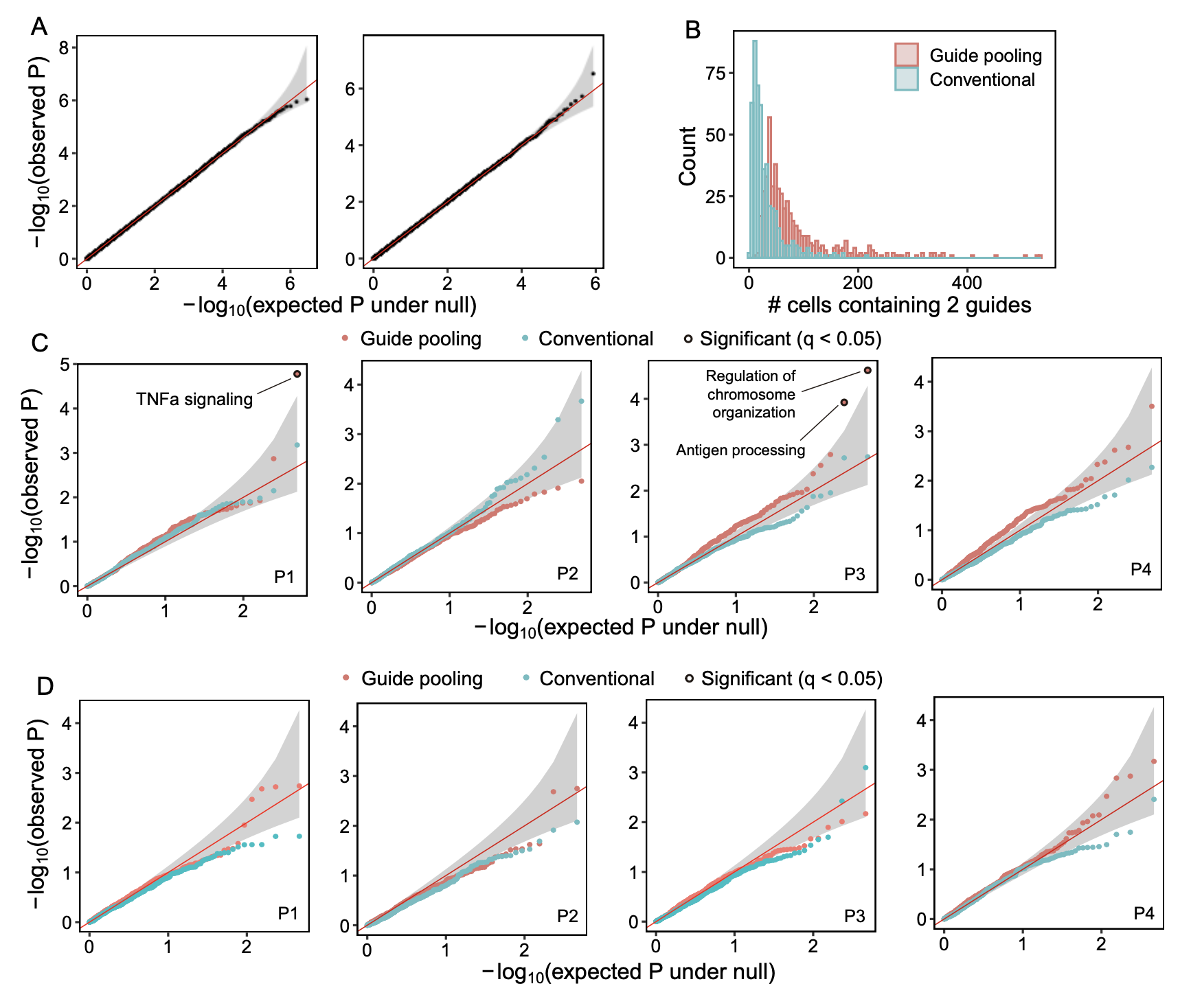

**Figure S5. Additional analyses of genetic interaction effects.** (**A**) Quantile-quantile plots comparing the distribution of observed interaction p-values for all perturbation pairs present in at least 5 cells on all downstream genes for the guide-pooled screen (left; 186 pairs * 16,268 downstream genes = 3,025,848 effects) and conventional screen (right; 46 pairs * 18,617 downstream genes = 856,382 effects) versus a uniform distribution under the null (x-axis). Grey region indicates the 95% confidence interval under the null. (**B**) Histogram of cells per co-functional modules (out of 490 total) containing at least 2 genes in the module. (**C**) Quantile-quantile plots comparing the distribution of observed intra-modular interaction p-values on the four gene programs (y-axis) versus a uniform distribution under the null (x-axis). Significant p-values (q < 0.05) are indicated with a black border and are labelled with the module name. (**D**) Same as **C**, but showing inter-modular (435 module pairs) rather than intra-modular interaction p-values.

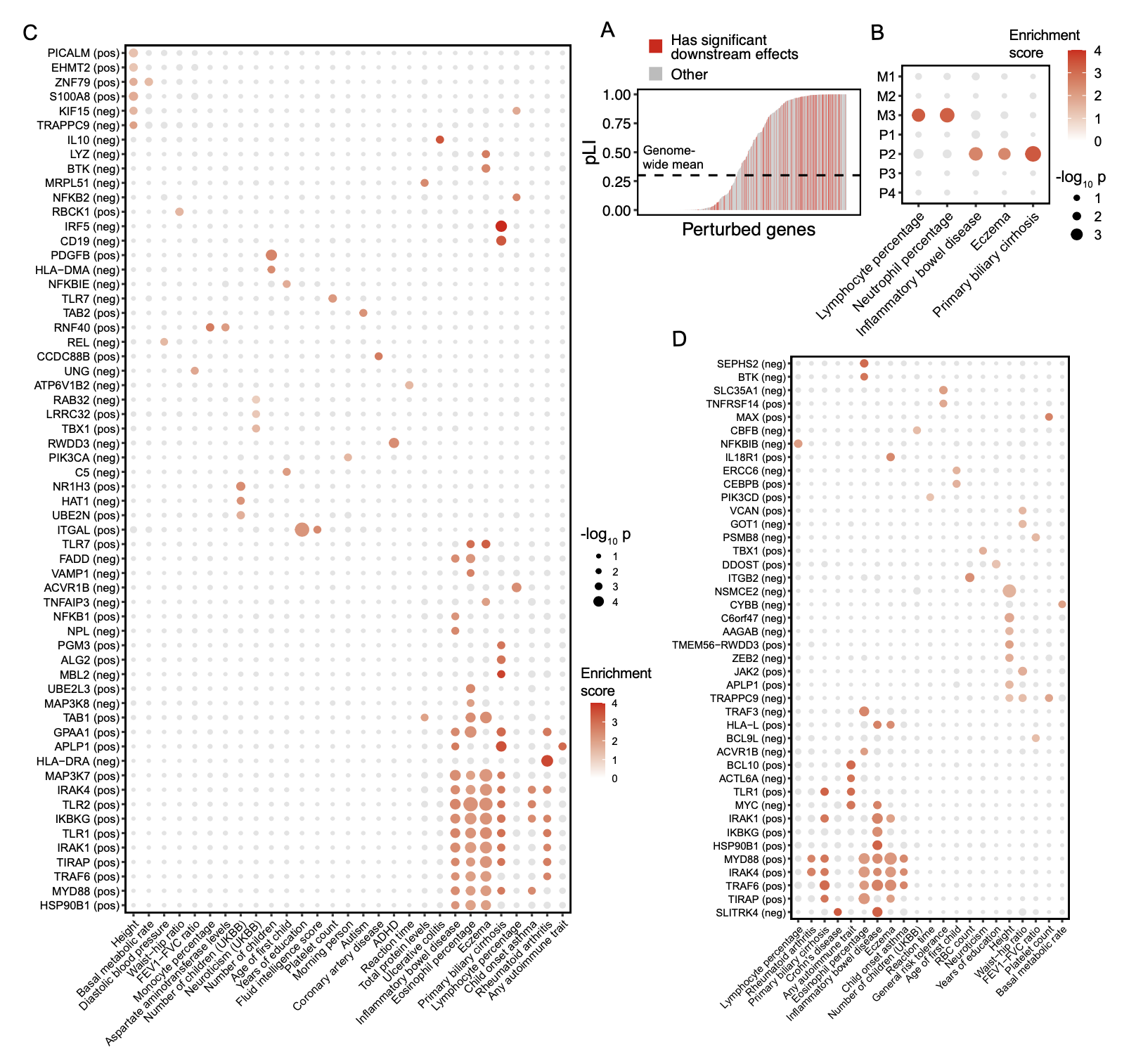

**Figure S6. Additional heritability analyses.** (**A**) Distribution of probability of loss of function intolerance (pLI; Karczewski et al., 2020; y-axis) of all 598 perturbed genes (x-axis). Perturbed genes with significant downstream effects (affecting the expression of >100 genes (q < 0.05) in either the knock-out or knock-down screen; N = 159) are indicated in red, with the remaining perturbed genes indicated in grey. Mean pLI across all genes is indicated with a dotted line. (**B**) Heritability enrichment scores (estimated using sc-linker; Jagadeesh et al., 2022) of perturbation modules M1-3 and gene programs P1-4. Shown are all traits with at least one significant (p < 0.001) effect (in red). (**C-D**) Heritability enrichment scores of all perturbation signatures from the knock-out (**C**) and knock-down (**D**) screens with at least one significant (p < 0.001) effect on any trait (in red).

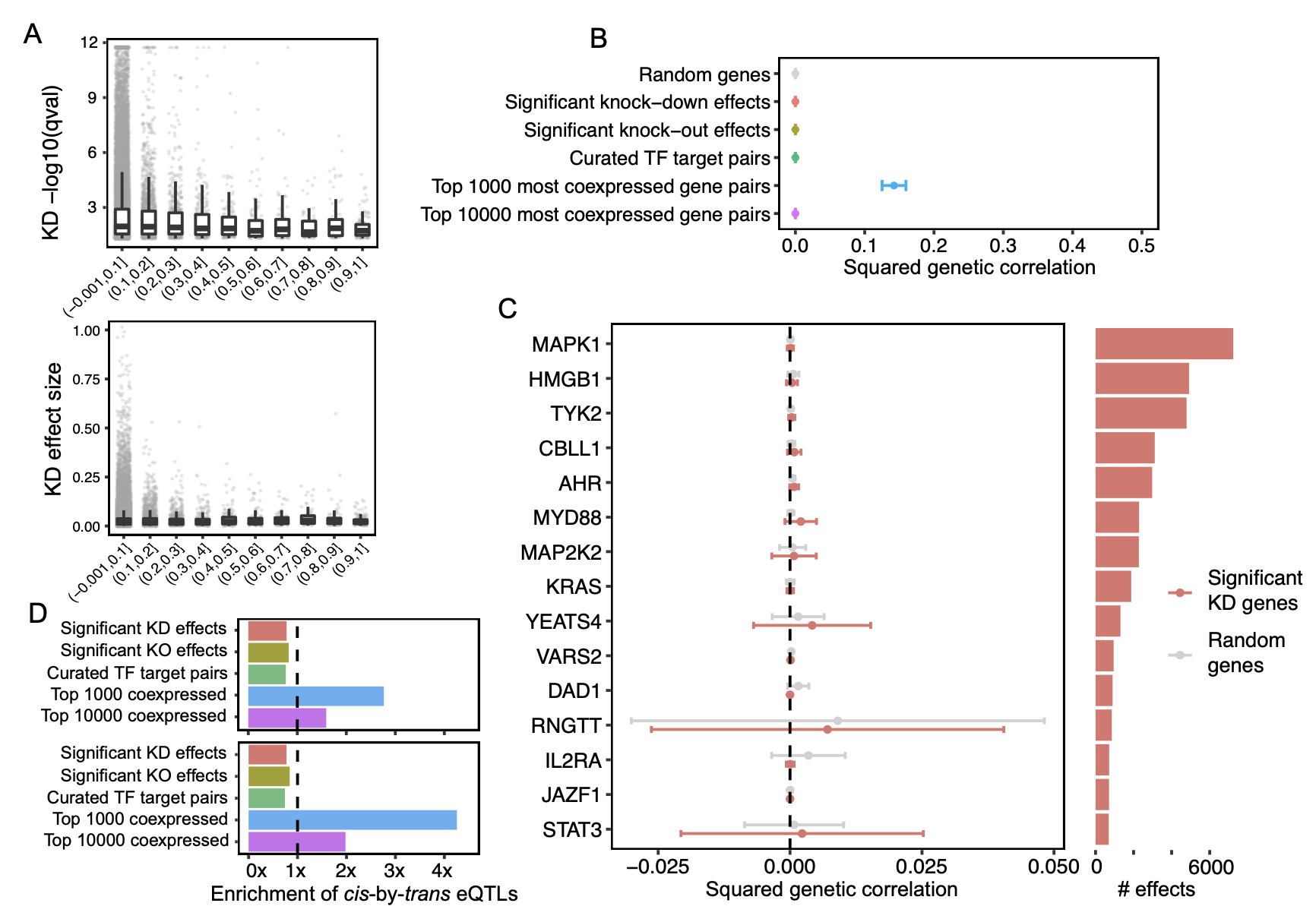

**Figure S7. Additional eQTL analyses.** (**A**) Relationship between knock-down significance (top) or effect size magnitude (bottom) versus probability of observing *cis*-by-*trans* eQTL in the Fairfax dataset (x-axis). (**B**) Genetic correlation estimates across various sources of gene-gene relationships. Estimates are meta-analyzed across all gene pairs. Error bars represent standard errors. (**C**) Genetic correlation estimates for significant gene-gene pairs from the knock-down screen stratified by perturbed gene. Barplot indicates the total number of significant downstream genes for each perturbed gene. (**D**) Same as **Figure 6D**, but varying the posterior probability threshold of determining a significant *cis*-by-*trans* eQTL to 0.25 (top) and 0.5 (bottom).

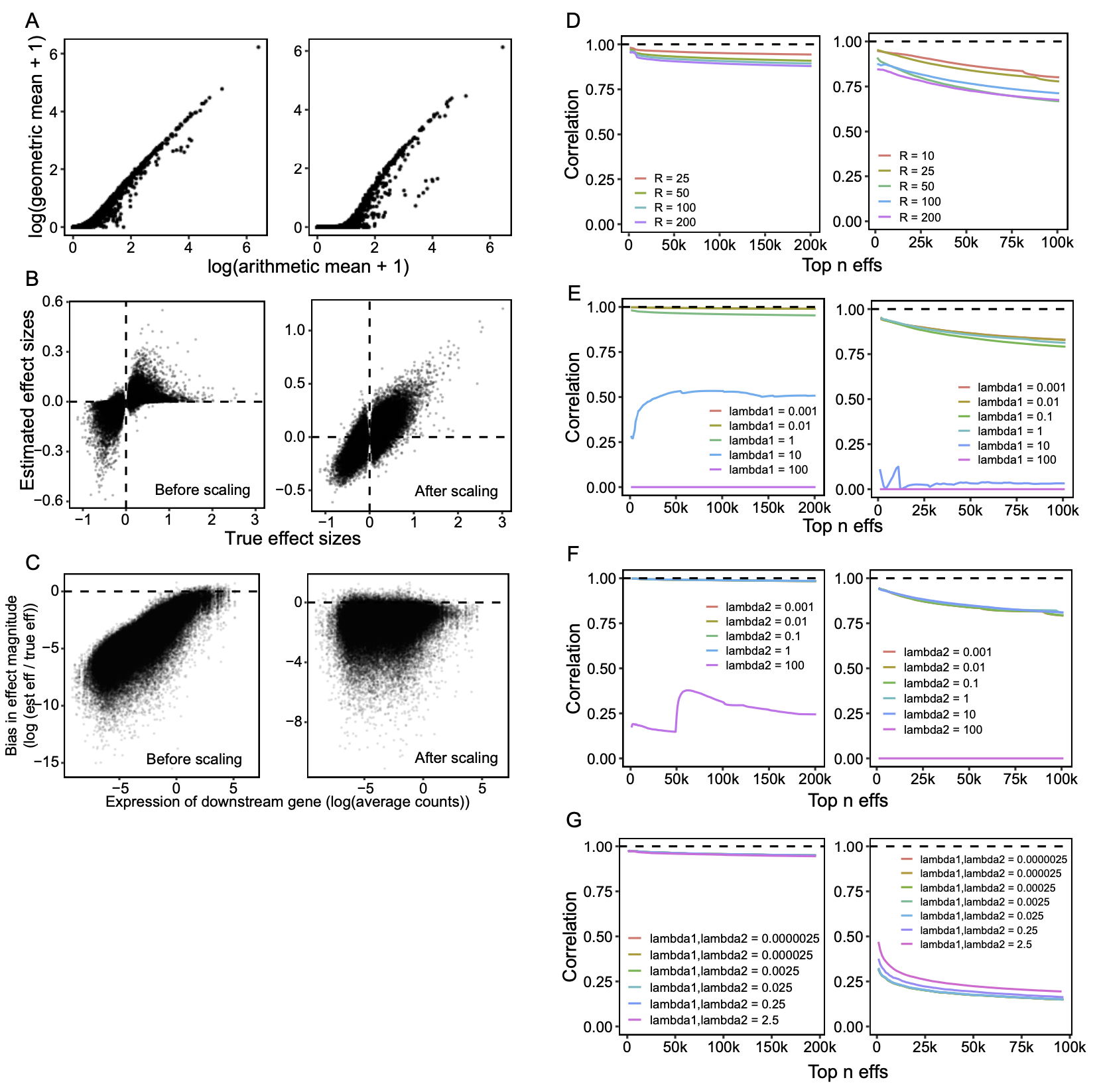

**Figure S8.** **Evaluating inference procedures.** (**A**) Arithmetic versus geometric mean of expression. Each point represents an individual gene’s mean expression across 3 (left) or 10 (right) random control cells from our conventional knock-out screen. (**B**) Impact of scaling effects from FR-Perturb in simulations. Scatterplots of true effect sizes (x-axis) versus estimated effect sizes (y-axis) in simulated cells before versus after scaling effects (**Methods**). True effect sizes were taken as the effects estimated from the knock-out screen, with 50,000 cells simulated. (**C**) Bias in effect size magnitude in estimated effects from **B** (y-axis) versus the expression level of the downstream gene (x-axis). (**D**) Impact of varying the hyperparameters in FR-Perturb. (Left) Correlation of top *n* most significant effects from the knock-out screen with the same effects estimated using the default FR-Perturb hyperparameters (R=10, lambda1=0.1, lambda2=10) when varying R. (Right) Correlation of top *n* most significant effects from the cell-pooled screen with the same effects from the conventional knock-out screen when varying R. (**E-F**) Same as **D**, but varying lambda1 (**E**) and lambda2 (**F**) rather than R. We note that the poor performance of large values of lambda1 and lambda2 is primarily due to most of the inferred effect sizes being zero. (**G**) Impact of varying hyperparameters of elastic net. (Left) Correlation of top *n* most significant effects from the knock-out screen with the same effects estimated using the default elastic net hyperparameters (lambda1=0.00025, lambda2=0.00025) when varying lambda1 and lambda2. (Right) Correlation of top *n* most significant effects from the cell-pooled screen with the same effects from the conventional knock-out screen when varying lambda1 and lambda2.

**Supplementary Note**

**Why does cell pooling reduce per-droplet signal?**

When matching the droplet count between our cell-pooled and conventional knock-out screens, we observed that cell pooling performs worse (in terms of out-of-sample validation accuracy) than the conventional screen (**Figure S3A**). This appears to contradict the notion that generating linear combinations of expression should increase per-measurement signal versus directly measuring expression counts [S1], but in fact is not entirely unexpected. In this section, we explain why this result occurs.

Consider a simple example where gene $i$ has an average expression of 100 counts (at whatever constant read depth) in unperturbed cells. Now consider three guides $a$, $b$, and $c$ which have a true log2-fold change effect of 1, 0, and 0 on gene $i$ respectively. Three cells containing each of guides $a$, $b$, and $c$ will have expression values of 200, 100, and 100 for gene $i$ respectively (in expectation). If we put these three cells in a droplet and measure their average expression, we will observe 133 counts in expectation, which is substantially closer to 100 than the 200 counts for the cell containing guide $a$. Moreover, including additional cells that do not have an effect on gene $i$ in the same droplet will further cause the average to be closer to 100 and thus decrease the signal to noise ratio, an effect that can be partially offset through deeper sequencing, which incurs additional cost. (As an aside, we note that with some forms of measurement, such as fluorescent measurements, but not sequencing-based counts, it may be possible to decrease the noise without increasing costs). On the contrary, for single cells containing all guides $a$, $b$, and $c$, assuming that the effect sizes of multiple guides combine additively in log expression space in the same cell, we will still observe an expression value of 200 for the cell. Adding more guides per cell will not dilute the signal.

Although it is not obvious that the effect size dilution in the cell pooling scenario will overpower the efficiency gains from measuring linear combinations of expression, in practice and in simulations we observed it to be the case (**Figure S3A**). In other words, when performing cell pooling, we are increasing the sample size per droplet by measuring multiple cells per droplet but decreasing the signal to noise ratio, and the latter ends up decreasing the overall per-droplet efficiency of cell pooling.

To summarize, the fact that cell pooling takes linear combinations of expression values across different cells (in which the presence of additional control cells will dilute the overall signal), whereas guide pooling takes linear combinations of effect sizes within the same cell (in which the presence of additional control guides will not dilute the overall signal), causes the performance of cell pooling (but not guide pooling) to drop when increasing the level of overloading.

**Use of guide pooling to systematically study high-order genetic interaction effects**

One major challenge in learning second-order interaction effects (and even more so for higher-order interaction effects) is the very large space of possible gene combinations that must be tested, wherein our guide-pooled approach can in theory lead to exponential increases in efficiency over the conventional approach. We describe these theoretical efficiency gains below.

For simplicity, suppose that we are measuring a scalar phenotype. If $N$ is the number of distinct perturbations and $W$ the order of the interaction effect among the perturbations, then there are $\binom{N}{W}$ total interaction effects to learn. Using the conventional approach, which involves individually perturbing each gene combination in each sample, one needs $O\left( \binom{N}{W} \right)\approx O\left( N^{W} \right)$ samples to learn all interaction effects. This quantity scales exponentially with respect to $W$ and polynomially (with exponent $W$) with respect to $N$. On the other hand, the guide-pooled approach involves perturbing multiple random gene combinations in each sample (cell). This requires only $O\left( k\log\binom{N}{W} \right)\approx O\left( k\log N^{W} \right)\approx O\left( Wk\log N \right)$ samples. This quantity scales *linearly* with respect to $W$ and logarithmically with respect to $N$, making it potentially tractable to systematically study genetic interaction effects at high order.

While the quantity $k$ (which refers to the number of nonzero interaction effects) will also increase with both $N$ and $W$, it is likely that $k$ will increase much more slowly than $\binom{N}{W}$, as supported by the observation that interaction effects get substantially sparser when going from second to third order interaction effects in yeast [S2]. When the goal is to study an $M$-dimensional phenotype like effect sizes on gene expression$, k$ in the above expression can be replaced by $r+q$ (where $r$ refers to the rank of the $\binom{N}{W}\times M$ interaction effect size matrix and $q$ refers to number of nonzero interaction effects on latent factors of the interaction effect size matrix). Essentially nothing is known about the low-rank structure of genetic interaction effects on gene expression profiles in any model system, though our analyses of inter- and intra-module interaction effects (**Figure 5F, S5C,D**) provides some initial evidence for the existence of such low-rank structure in second-order interaction effects.

Importantly, these efficiency gains are also contingent on the fact that the additive model holds across all orders. This means, for example, that the phenotype of a sample (cell) containing perturbations for genes $i$, $j$, and $k$ should on average be the sum of all first order effects as well as all *pairs* of second order effects between the three perturbations. In practice, very little is known about the structure of higher-order interaction effects and how they systematically combine within a cell, so more study is required to investigate the validity of this model.

**Enrichment of *cis*-by-*trans* eQTLs in the most co-expressed genes suggests the presence of confounding**

In **Figure 6D**, we observed that *cis*-by-*trans* eQTLs are enriched in gene pairs with the highest correlation of expression across samples (*i.e.*, the most co-expressed genes). In this section, we explain why this observation suggests the presence of confounders in the eQTL dataset, even though we followed established protocols to remove them in our analyses (*i.e.*, through the inclusion of top expression principal components).

Under random mating, genetic effects are independent of environmental effects, which enables eQTL effects to be viewed as perturbations on gene expression. However, gene co-expression is almost completely dominated by environmental rather than genetic effects [S3], so the fact that we observe an enrichment of *cis*-by-*trans* eQTLs in the most co-expressed genes (**Figure 6D**) is evidence of an unexpected relationship between genetic correlation and environmental correlation (note that this can occur even if genetic and environmental effects are independent, see below). There are many possible causal scenarios that can lead to a relationship between these quantities. We illustrate three of the simplest scenarios below:

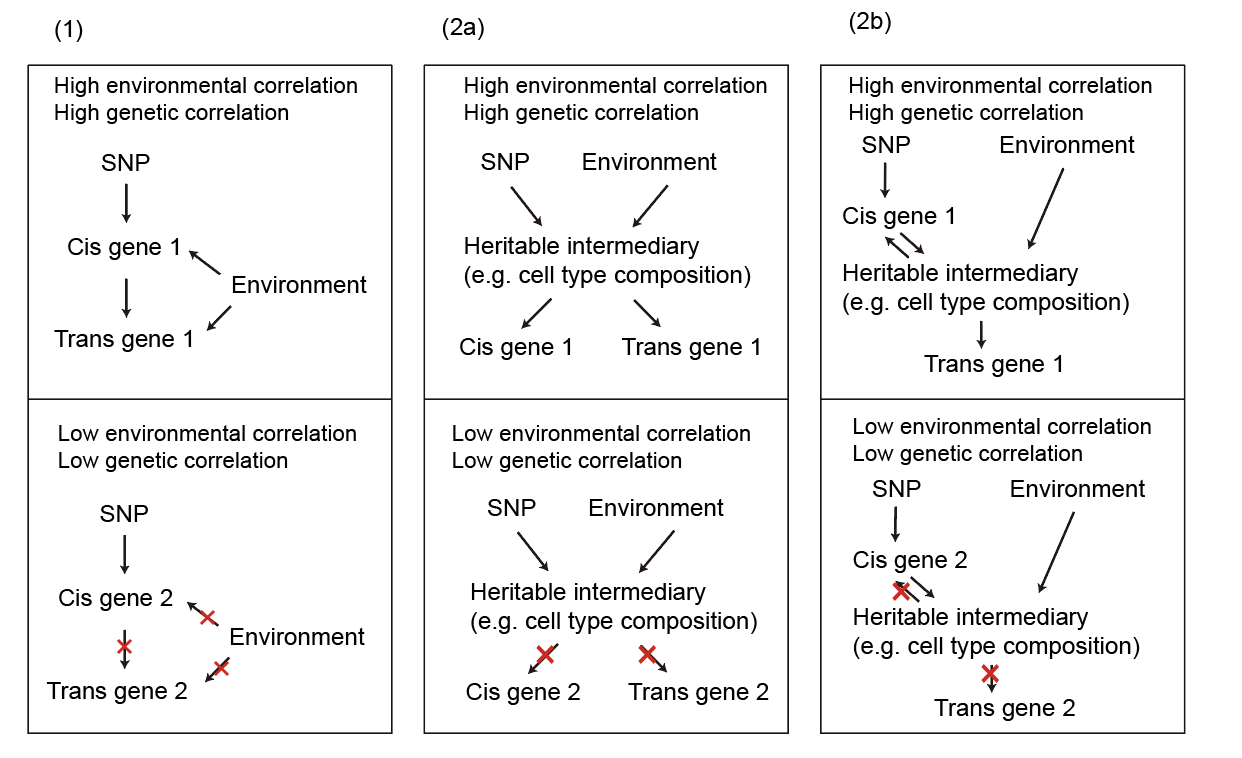

Each column in the above figure corresponds to a single scenario applied to different gene pairs (top and bottom boxes). The top box illustrates the scenario applied to a gene pair with high genetic and environmental correlation, while the bottom box illustrates the same scenario applied to a different gene pair with low genetic/environmental correlation. Scenarios (1) and (2b) have a causal link from the *cis* gene to the *trans* gene, but scenario (2a) does not and is driven by confounding. Although all three scenarios are possible in theory, we discuss below why scenario (2a) is probably the most common one.

Scenario (1) requires that the probability of the *cis* gene having a causal effect on the *trans* gene is related to the environmental correlation between the genes. Although this can in theory be explained by various biologically plausible scenarios (*e.g.*, the *cis* gene only affects the *trans* gene in the presence of a co-factor which is influenced by the environment), this scenario is less parsimonious in light of our observation that *cis*-by-*trans* eQTLs are much more strongly enriched in the top 1,000 most co-expressed gene pairs than the top 10,000 most co-expressed gene pairs (**Figure 6D**). In other words, this would mean that the hypothetical environmentally-driven co-factor scenario acts much more strongly on the top 0.0003% of gene pairs than the top 0.003% of gene pairs, both of which are a tiny and somewhat arbitrary (from the perspective of biological function) fraction of all possible gene pairs. For the top 0.0003% of gene pairs to be functionally different than the top 0.003% when the degree of co-expression is similar between the two groups, an extremely (and we argue unrealistically) non-linear relationship between genetic and environmental correlation would need to be present.

Meanwhile, scenarios (2a) and (2b) involve the presence of a heritable intermediary between the SNP and the *cis* and *trans* genes which can also be affected by the environment. This intermediary could be intercellular/cell-type heterogeneity (which is known to be a heritable and polygenic trait in blood cells [S4]) or a molecular pathway. The intermediary causes some genes’ expression to be more correlated with each other than others. Thus, because gene co-expression is determined by the intermediary and both genetic and environmental factors flow through it, the genes pairs with the highest environmental correlation and genetic correlation are the same gene pairs. This would explain why we observe significantly stronger enrichment of *cis*-by-*trans* eQTLs in the top 1,000 gene pairs than the top 10,000 gene pairs. Between scenarios (2a) and (2b), (2b) requires a bi-directional causal effect between the *cis* gene and the intermediary, so (2a) is more parsimonious.

We note that the presence of substantial intercellular heterogeneity between individuals is well-established in eQTL studies [S5], even those in relatively homogenous cell specimens. For example, in Fairfax et al. [S6], the inclusion of gene expression principal components to account for intercellular heterogeneity (despite the cells being the same cell type) results in an increase in the number of *cis*-eQTLs detected (from ~4500 to ~7200) and a decrease in the number of *trans*-eQTLs (from ~700 to ~350) (Fig. S2 in Fairfax et al.). This pattern is consistent with the observation that confounding (due to intercellular heterogeneity and/or other factors) decreases the number of detected cis-eQTLs while increasing the number of false-positive *trans*-eQTLs [S7], in which case controlling for confounding would reverse these patterns as observed in Fairfax et al. However, we note that the observed decrease in detected *trans*-eQTLs when controlling for gene expression principal components can in theory also be explained by principal components capturing true, widespread intra-cellular *trans* effects. Disentangling spurious inter-cellular effects from true intra-cellular *trans* effects remains a key challenge in *trans*-eQTL studies.

**Explaining depletion of *cis*-by-*trans* eQTLs among regulatory gene pairs from Perturb-seq**

We observed that regulatory gene pairs identified by our Perturb-seq screens are substantially depleted for colocalized *cis*-by-*trans* eQTLs compared to random gene pairs (**Figure 6D**). In the scenario that Perturb-seq does not capture regulatory relationships in the eQTL dataset, we might expect that Perturb-seq gene pairs are essentially a random selection of genes, and thus we would observe the *same* probability of colocalization of *cis*-by-*trans* eQTL among our Perturb-seq gene pairs versus random gene pairs. However, the fact that we observe a *lower* probability of colocalization of *cis*-by-*trans* eQTLs among Perturb-seq gene pairs versus random gene pairs requires an additional explanation. Two potential explanations are discussed below.

(1) *Perturb-seq gene pairs are less confounded than random gene pairs.* We have previously illustrated that detection of *cis*-by-*trans* eQTLs is likely to be at least partially driven by confounders such as intercellular heterogeneity (see previous section). We have additionally observed that significant Perturb-seq gene pairs are more likely to be under selective constraint than random genes, and that highly co-expressed genes are less likely to be under selective constraint (**Figure 6E**). Thus, under a model where *cis*-by-*trans* eQTLs are driven by confounding, we expect to see an enrichment of *cis*-by-*trans* eQTLs for co-expressed (and unconstrained) gene pairs, and a concomitant depletion of *cis*-by-*trans* eQTLs for significant Perturb-seq (and constrained) gene pairs. The observed depletion can thus be explained by a negative relationship between the technical confounding and selective constraint, in the presence of *cis*-by-*trans* eQTLs driven by confounding.

(2) *Negative selection on the trans gene in Perturb-seq gene pairs reduces power to detect context-specific cis-by-trans eQTLs compared to random gene pairs*. We observed that *trans* genes in the Perturb-seq gene pairs are under more selective constraint than average genes (**Figure 6E**). However, under the simplest model of SNPs acting on the *trans* gene through the *cis* gene (see below), this fact alone does not manifest in decreased power to detect *cis*-by-*trans* eQTLs compared to random gene pairs (because our analysis explicitly conditioned on the *cis*/perturbed gene when selecting random gene pairs).

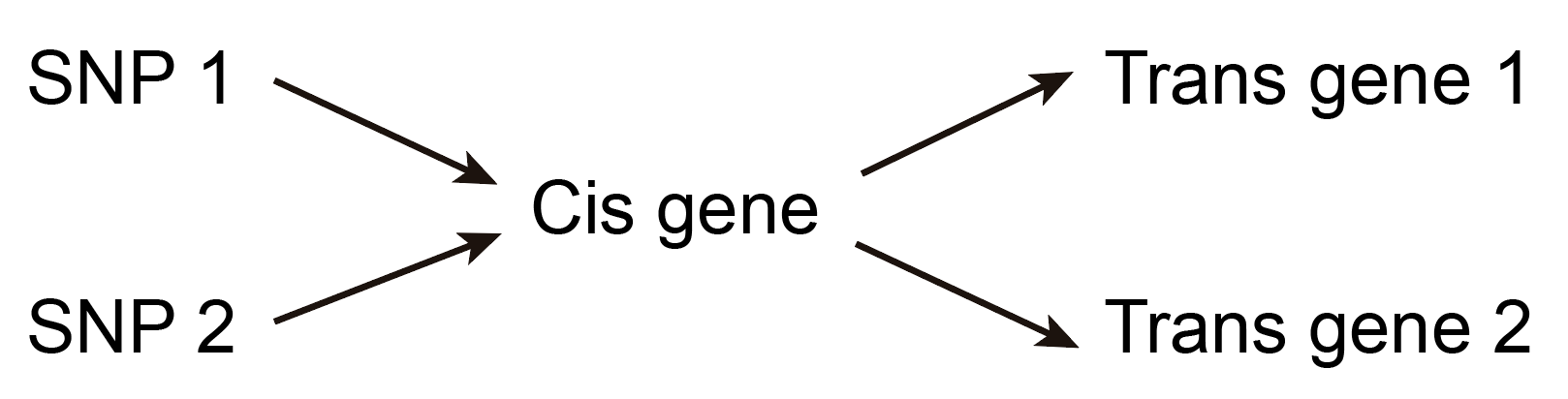

In this model, every SNP that affects the *cis* gene (including SNPs 1 and 2) will also affect both *trans* genes. Suppose that *trans* gene 1 is under strong selective constraint, so that any SNP with an effect on it is selected out of the population, whereas *trans* gene 2 is not under any selective constraint. All SNPs that regulate the *cis* gene (including both SNP 1 and SNP 2) will have decreased frequency in the population, resulting in uniformly decreased power to detect *cis*-by-*trans* eQTLs for both *cis* gene -> *trans* gene 1 and *cis* gene -> *trans* gene 2 (as well as any additional *trans* genes that the *cis* gene regulates). As noted above, when we compared Perturb-seq gene pairs to random gene pairs in our analyses, we kept the *cis* gene constant in all comparisons, thus controlling for differences in power for SNPs that regulate the *cis* gene.

A slightly modified version of the above model is needed to explain our observation of a depletion of *cis*-by-*trans* eQTLs for Perturb-seq gene pairs:

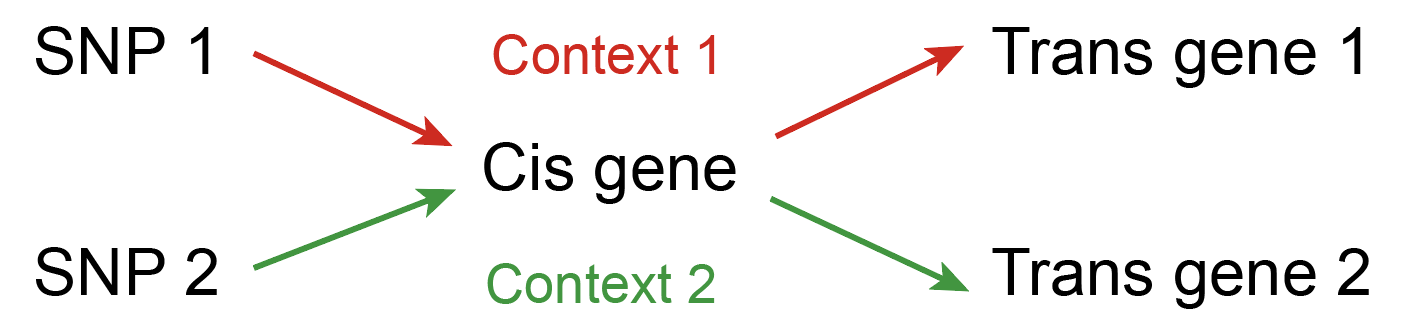

Here, SNP 1 only acts on *trans* gene 1 through the *cis* gene in a given context 1 (which could be a cell or tissue type, transient cell state, intracellular location, etc.), while SNP 2 only acts on *trans* gene 2 through the *cis* gene in a different context 2. Under this model, selection on *trans* gene 1 causes SNP 1 to be selected out of the population but not SNP 2. This will decrease the power to detect *cis*-by-*trans* eQTLs for *cis* gene -> *trans* gene 1 but not *cis* gene -> *trans* gene 2. Thus, differential selection on context-specific *cis*-eQTLs for the same gene (as has been previously hypothesized [S8] could explain why we observe a depletion of *cis*-by-*trans* eQTLs for *trans* genes under selection.
